# Folding of Prestin’s Anion-Binding Site and the Mechanism of Outer Hair Cell Electromotility

**DOI:** 10.1101/2023.02.27.530320

**Authors:** Xiaoxuan Lin, Patrick Haller, Navid Bavi, Nabil Faruk, Eduardo Perozo, Tobin R. Sosnick

**Affiliations:** Department of Biochemistry and Molecular Biology, The University of Chicago, Chicago, Illinois, USA; Center for Mechanical Excitability, The University of Chicago, Chicago, Illinois, USA; Institute for Neuroscience, The University of Chicago, Chicago, Illinois, USA; Institute for Biophysical Dynamics, The University of Chicago, Chicago, Illinois, USA; Prizker School for Molecular Engineering, The University of Chicago, Chicago, Illinois, USA

**Keywords:** Mass spectrometry, hydrogen exchange, cochlear amplification, electromotility

## Abstract

Prestin responds to transmembrane voltage fluctuations by changing its cross-sectional area, a process underlying the electromotility of outer hair cells and cochlear amplification. Prestin belongs to the SLC26 family of anion transporters yet is the only member capable of displaying electromotility. Prestin’s voltage-dependent conformational changes are driven by the putative displacement of residue R399 and a set of sparse charged residues within the transmembrane domain, following the binding of a Cl^-^ anion at a conserved binding site formed by amino termini of the TM3 and TM10 helices. However, a major conundrum arises as to how an anion that binds in proximity to a positive charge (R399), can promote the voltage sensitivity of prestin. Using hydrogen-deuterium exchange mass spectrometry, we find that prestin displays an unstable anion-binding site, where folding of the amino termini of TM3 and TM10 is coupled to Cl^-^ binding. This event shortens the TM3-TM10 electrostatic gap, thereby connecting the two helices, resulting in reduced cross-sectional area. These folding events upon anion-binding are absent in SLC26A9, a non-electromotile transporter closely related to prestin. Dynamics of prestin embedded in a lipid bilayer closely match that in detergent micelle, except for a destabilized lipid-facing helix TM6 that is critical to prestin’s mechanical expansion. We observe helix fraying at prestin’s anion-binding site but cooperative unfolding of multiple lipid-facing helices, features that may promote prestin’s fast electromechanical rearrangements. These results highlight a novel role of the folding equilibrium of the anion-binding site, and helps define prestin’s unique voltage-sensing mechanism and electromotility.

## Introduction

Hearing sensitivity in mammals is sharply tuned by a cochlear amplifier associated with electromotile length changes in outer hair cells^1^. These changes are driven by prestin (SLC26A5), a member of the SLC26 anion transporter family, which converts voltage-dependent conformational transitions into cross-sectional area changes, affecting its footprint in the lipid bilayer^2^. This process plays a major role in mammalian cochlear amplification and frequency selectivity, with prestin knockout producing a 40-60 dB signal loss in live cochleae^3^. Unlike most molecular motors, where force is exerted from chemical energy transduction, prestin behaves as a putative piezoelectric device, where mechanical and electrical transduction are coupled^4^. As a result, prestin functions as a direct voltage-to-force transducer. Prestin’s piezoelectric properties are unique among members of the SLC26 family, where most function as anion transporters.

Recent structures determined by cryo-electron microscopy (cryo-EM) have sampled prestin’s conformational space under various anionic environments and located the anion-binding site at the electrostatic gap between the amino termini of TM3 and TM10 helices^5–7^. This anion-binding pocket is highly conserved, and is influenced by surrounding hydrophobic residues in TM1 and by a fixed positive charge from residue R399 on TM10. Movements of this binding site are coupled to the complex reorientation of the core domain relative to the gate domain^5,6^, reminiscent of the conformational transitions in transporters displaying an elevator-like mechanism^8^. Prestin exhibits minimal transporter ability yet is structurally similar to the non-electromotive anion transporter SLC26A9 (sequence identity = 34%; Cα RMSD = 3.4 Å for the transmembrane domain (TMD), PDB: 7S8X and 6RTC)^7,9,10^. Questions remain as to the molecular basis underlying the distinct functions of the two proteins. Importantly, the role of bound anions, which is required for prestin electromotility^11^, is still elusive.

Prestin’s voltage dependence is tightly regulated by intracellular anions of varying valence and structure^12,13^, whereas anion affinity is also regulated by voltage and tension^14^. These phenomena suggest that anions, rather than behaving as explicit gating charges, may serve as allosteric modulators^14^. Incorporating a fixed charge alternative to a bound anion through an S398E mutation preserves prestin’s nonlinear capacitance (NLC) but results in insensitivity to salicylate, a strong competing anionic binder^7^. Except for residue R399, charged residues located in the TMD distribute towards the membrane-water interface^5–7^ and display minimal contributions to the total gating charge estimated from NLCs^15^. Electrostatic calculations show that R399 has a strong contribution to the local electrostatics at the anion-binding site, by providing ∼40% of the positive charge at the bilayer mid-plane^5^. However, the existing structural and functional data cannot explain why prestin’s voltage dependence requires close proximity of both a negative charge (the bound anion or S398E)^11^ and a positive charge (R399^5,16^). The resolution of this conundrum will define an essential step towards our understanding of prestin’s unique voltage-sensing mechanism.

Here we studied the influence of anion-binding on the dynamics and structural changes of prestin as a function of anions (Cl^-^, SO_4_^2-^, salicylate, and HEPES) via hydrogen-deuterium exchange mass spectrometry (HDX-MS). The HEPES condition was achieved by Cl^-^ removal (dialysis), which inhibits prestin’s NLC^11^. In a HEPES-based buffer, prestin’s NLC shifts to depolarized potentials, associated to a more expanded state at 0 mV that is coupled to low anion affinity^12,13^. Based on the above studies and the large size of HEPES anions, we assumed minimal binding of HEPES anions to prestin and hence associated HEPES condition to a putative apo state in this study. By comparing the dynamics of prestin with its close non-piezoelectric relative, the anion transporter SLC26A9, we identified distinct features unique to prestin, including a relatively unstable anion-binding site that folds upon binding, thereby allosterically modulating the dynamics of the TMD. In contrast, the stability and hydrogen-bond pattern of SLC26A9’s anion-binding site were minimally affected by anion binding, albeit displaying high similarities to prestin in both structure and sequence. Prestin reconstituted in nanodisc exhibited indistinguishable dynamics compared to detergent-solubilized prestin, except for a destabilized TM6 which mediates prestin’s mechanical expansion^5^. We observed fraying of the helices involved in the binding site whereas cooperative unfolding of multiple lipid-facing helices including TM6, which may explain prestin’s fast and large-scale motions. These results highlight the significance of the anion-binding site’s folding equilibrium in defining the unique properties of prestin’s voltage dependence and electromotility.

## Results

We carried out HDX measurements on dolphin prestin and mouse SLC26A9 solubilized in glyco-diosgenin (GDN) at either pD_read_ 7.1, 25 °C or pD_read_ 6.1, 0 °C (**Table S1**). The observed HDX rates reported on the stability, as exchange occurred mostly via EX2 kinetics (**Supporting Information Text 1**). Employment of the two conditions increased the effective dynamic range of the HDX measurement to span seven log units, allowing us to determine the stability of both the highly and minimally stable regions within the protein^17^. To properly combine the two datasets, the stability of the protein should be the same under the two conditions, and this was supported by the exchange rates scaling with *k*_*chem*_ (intrinsic exchange rates)^18,19^ (**Fig S7A**).

The HDX data were presented in terms of the relevant region with the specific sequence and peptides noted in parentheses (**Materials and Methods**), e.g., the N-terminus of prestin’s TM10 (Region_394–397_: Peptide_392–397_). Although we mostly focused on the anion-binding site, we also obtained comparative thermodynamic information throughout the two proteins (**Supporting Information Text 2**).

### Prestin’s anion-binding site is less stable than SLC26A9’s

To examine the effect of anion binding to the dynamics of prestin and SLC26A9, we dialyzed the proteins purified in Cl^-^ into a HEPES buffer lacking other anions. Cl^-^ removal resulted in distinct stability changes for prestin and SLC26A9, manifested by significant HDX acceleration for prestin while mild HDX slowing for SLC26A9 (**Fig. 1 & Fig. S1**). These HDX effects indicate that anion binding induced global stabilization for prestin while slight destabilization for SLC26A9 (**Fig. 1**).

**Figure 1:**
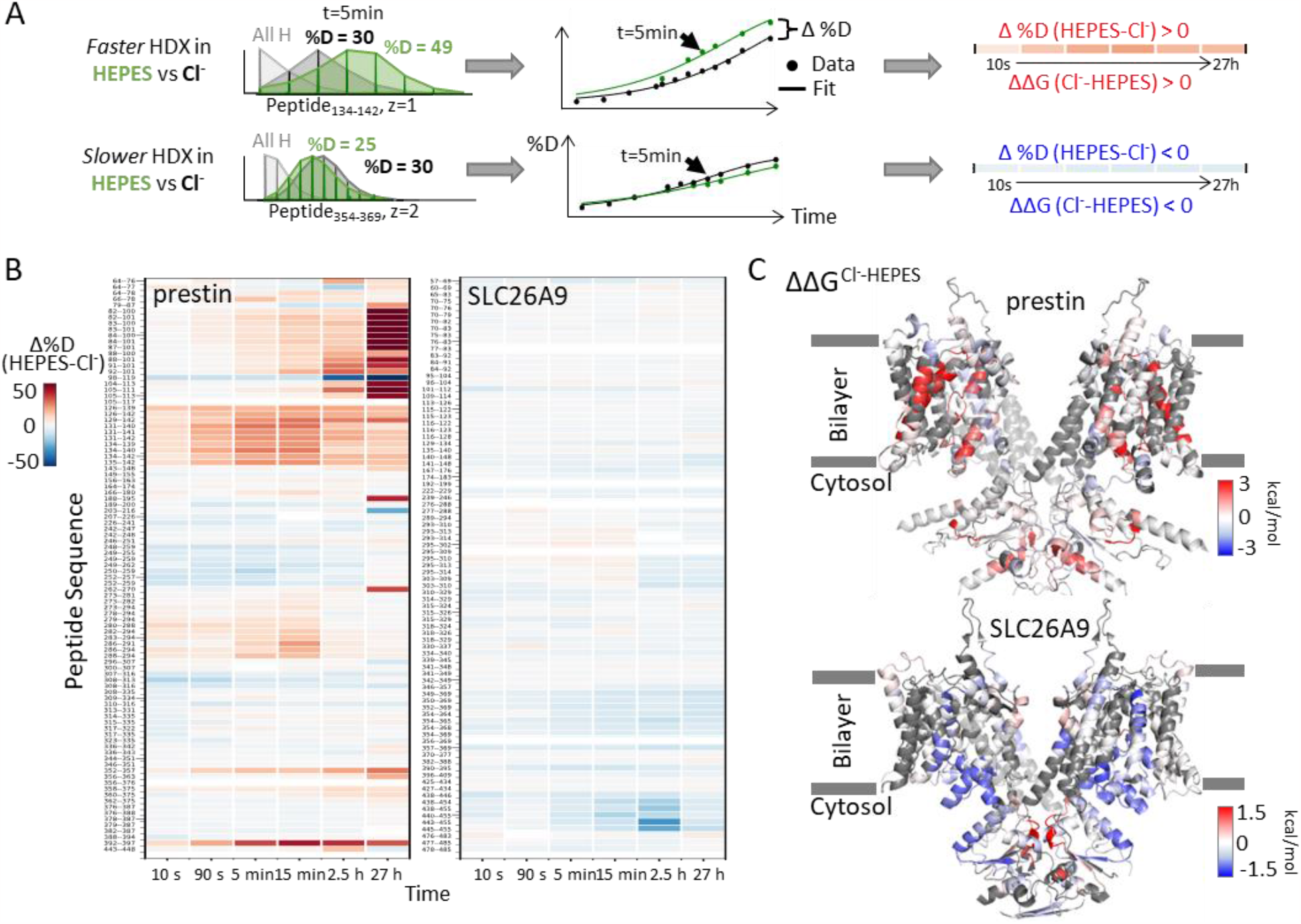
Distinct HDX response of prestin and SLC26A9 to Cl^-^ binding. **(A)** HDX data analysis to obtain **(B)** and **(C)**. One example peptide is shown in cases where HDX becomes faster or slower in HEPES (the putative apo state) compared to in Cl^-^. Deuteration levels are obtained from the mass spectra. Here spectra for the undeuterated peptide (grey) and after 5 min HDX labeling in Cl^-^ (black) and HEPES (green) are shown as an example. The resulting deuterium uptake plots are used to generate the differential deuteration heatmaps in **(B)**. Changes in free energy of unfolding (ΔΔG) in are calculated after fitting the data with a stretched exponential (**Materials and Methods**)^17^. **(B)** Heatmaps showing the difference in deuteration levels at each labeling time for all TMD peptides of prestin and SLC26A9 measured in HEPES compared to Cl^-^. Peptide sequences are displayed on the y-scale and legible through the high resolution image. **(C)** The ΔΔGs in HEPES compared to Cl^-^ for full-length prestin and SLC26A9 mapped onto the structure (PDB 7S8X and 6RTC). Red and blue indicate increased and decreased stability upon Cl^-^ binding, respectively. Following regions of the left subunits are shown as low transparency to highlight the binding site – prestin: TM5 and TM12-14; SLC26A9: TM5 and TM13-14. Regions with no fitting results are in grey.

Among the observed HDX responses for prestin, the HDX acceleration at the anion-binding pocket appeared to be the most pronounced and indicates local stabilization induced by anion binding (**Fig. 2A**). In detail, HDX accelerated by 20-fold for the N-termini of both TM3 (Region_136-142_: Peptide_134-142_ + 9 other peptides) and TM10 (Region_394–397_: Peptide_392–397_) (**Fig. 2A & Fig. S9.22-9.31**). This HDX change translates to a difference in free energy of unfolding (∆∆G) by at least 1.8 kcal/mol; 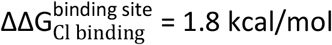. At least four residues in the middle of TM1 exhibited faster HDX (Region_90-101_: Peptide_88-101_ + 10 other peptides), collectively by 350-fold;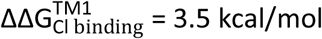 (**Fig. 1A & Fig. S9.6-9.16**). The TM1 region with accelerated HDX included L93, Q97, and F101, residues that are known to participate in the binding pocket^5,6,20^.

**Figure 2:**
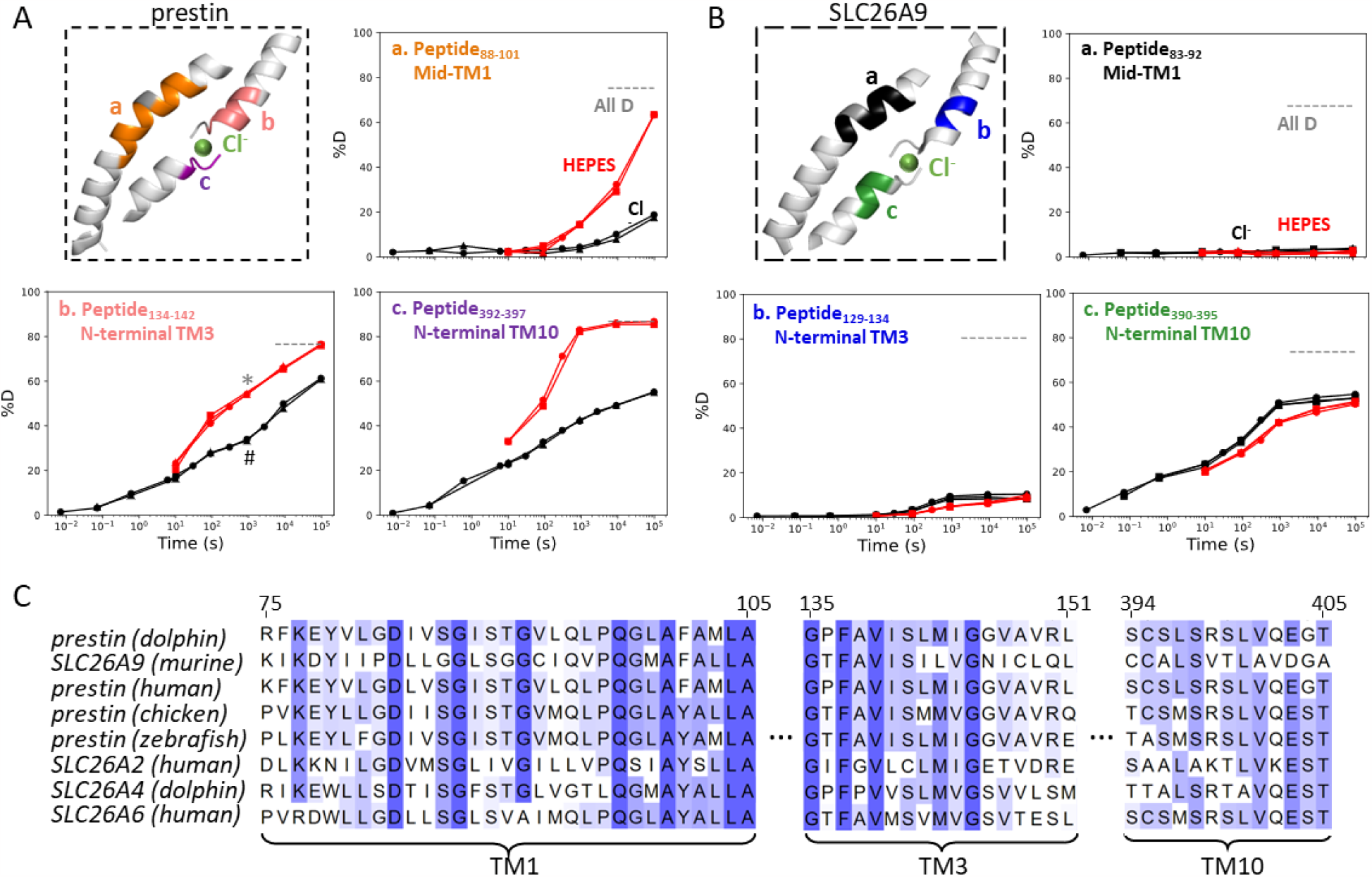
The anion-binding pockets for prestin and SLC26A9 exhibit distinct stability changes upon Cl^-^ binding, albeit highly conserved. **(A-B)** Cl^-^ binding stabilizes prestin’s anion-binding pocket **(A)** but mildly affects SLC26A9’s **(B)**. The structure shows the anion-binding pocket (TM1, TM3, and TM10) with the putative position of the bound Cl^-^. Colored regions correspond to peptides whose deuterium uptake plots are shown when the protein is in Cl^-^ (black) and in HEPES (red). Prolines are colored in grey. Grey dashed lines indicate deuteration levels in the full-D control. Data from two and three biological replicates are shown for prestin in Cl^-^ and HEPES, respectively. Data from three technical replicates are shown for SLC26A9. Replicates are shown as circles, triangles, and squares. Some replicates are superimposable and hence not observable. The symbols (* and #) in **(A.b)** denote data points used in **Fig. 3B. (C)** Sequence alignment using Clustal Omega of prestin and close SLC26 transporters across species for the anion-binding pocket. Shades of blue indicate degree of conservation.

SLC26A9 exhibited similar stability as prestin in Cl^-^ for the majority of the TMD, except for the N-terminal TM3 (Region_131-134_: Peptide_129-134_) which exchanged at least 100-fold slower than that of prestin’s (Region_136-142_: Peptide_134-142_) (**Fig. 2**). This difference in HDX points to a relatively unstable anion-binding site of prestin as compared to SLC26A9; 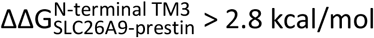 and was also seen in the site-resolved PFs that were obtained by deconvoluting the HDX-MS data using PyHDX^21^ (**Fig. S2**).

Compared to the 20∼350-fold HDX acceleration observed at prestin’s binding site upon Cl^-^ removal, HDX of SLC26A9’s binding pocket was only affected mildly (**Fig. 2B**). These included a slight slowing in HDX for the N-termini of TM3 (Region_131-134_: Peptide_129-134_) and TM10 (Region_392-395_: Peptide_390-395_) (**Fig. 2B**). The TM1 (Region_72-92_: Peptide_83-92_ + 10 other peptides) continued to remain undeuterated even after 27 h (**Fig. 2B & Fig. S10.4-10.14**), emphasizing the intrinsic high stability of SLC26A9’s anion-binding pocket.

Although the anion-binding pocket is highly conserved and structurally similar across members of the SLC26 family and SLC26A5 families (**Fig. 2C**), mammalian prestin is the only member capable of displaying eletromotility^22^. Hence, the distinct stability responses we observe for dolphin prestin and mouse SLC26A9 point to a prestin’s unique adaptation as a motor protein.

In addition to the binding pocket, we observed stability changes in various regions of the TMDs for prestin and SLC26A9 that may explain their distinct functions. For prestin, anion binding resulted in stabilization for the intracellular cavity but destabilization for regions facing the extracellular milieu (**Fig. 1C & Fig. S1A**). The stabilizing effects for the cytosol-facing regions were manifested by HDX acceleration upon Cl^-^ removal at the linker between TM2 and TM3, and the intracellular portions of TM7, TM8, & TM9 (Region_128-135_, Region_284-294_, & Region_354-375_) (**Fig. S9.22-9.27, S9.58-9.64, S9.82-9.87**). In contrast, HDX slowed for the regions facing the extracellular environment, namely the extracellular ends of TM5b, TM6, and TM7 (Region_250-262 &_ Region_309-316_) (**Fig. S9.45-9.51, S9.66-9.68, S9.71**). However, for SLC26A9, anion binding destabilized the cytosol-facing regions, as HDX slowed by ∼5-fold upon Cl^-^ removal for the intracellular ends of TM8, TM9, and TM12 (Region_351-369 &_ Region_440-455_) (**Fig. 1C, Fig. S1B, & Fig. S10.62-10.63, S10.77-10.82**). The distinct thermodynamic consequences of anion binding for prestin and SLC26A9 point to a distinct molecular basis underlying their different functions as a motor and a transporter, respectively.

### Anion binding drives the folding of prestin’s binding site

For prestin in HEPES, which adopted the putative apo state, the 20-fold HDX acceleration for the binding site (**Fig. 2A**) is consistent with a process of local destabilization, even unfolding, or increased solvent accessibility as the region becomes exposed to the intracellular water cavity^5^. To investigate these possibilities, we measured prestin’s HDX in response to a chaotrope, urea, which destabilizes proteins by interacting with backbone amides^23^. In a background of 360 mM Cl^-^, the addition of 4 M urea accelerated HDX for the N-terminus of TM3 (Region_137-140_: Peptide_134-140_) by 20-fold (**Fig. 3A**), suggesting that this region was destabilized and accessible to urea in its exchange-competent state. In apo prestin, however, the PF at the N-terminus of TM3 was unaffected by urea (after accounting for the known ∼50% slowing of the *Kchem*^23^) (**Fig. 3A**), arguing that this region was already unfolded prior to the addition of urea^24^.

**Figure 3:**
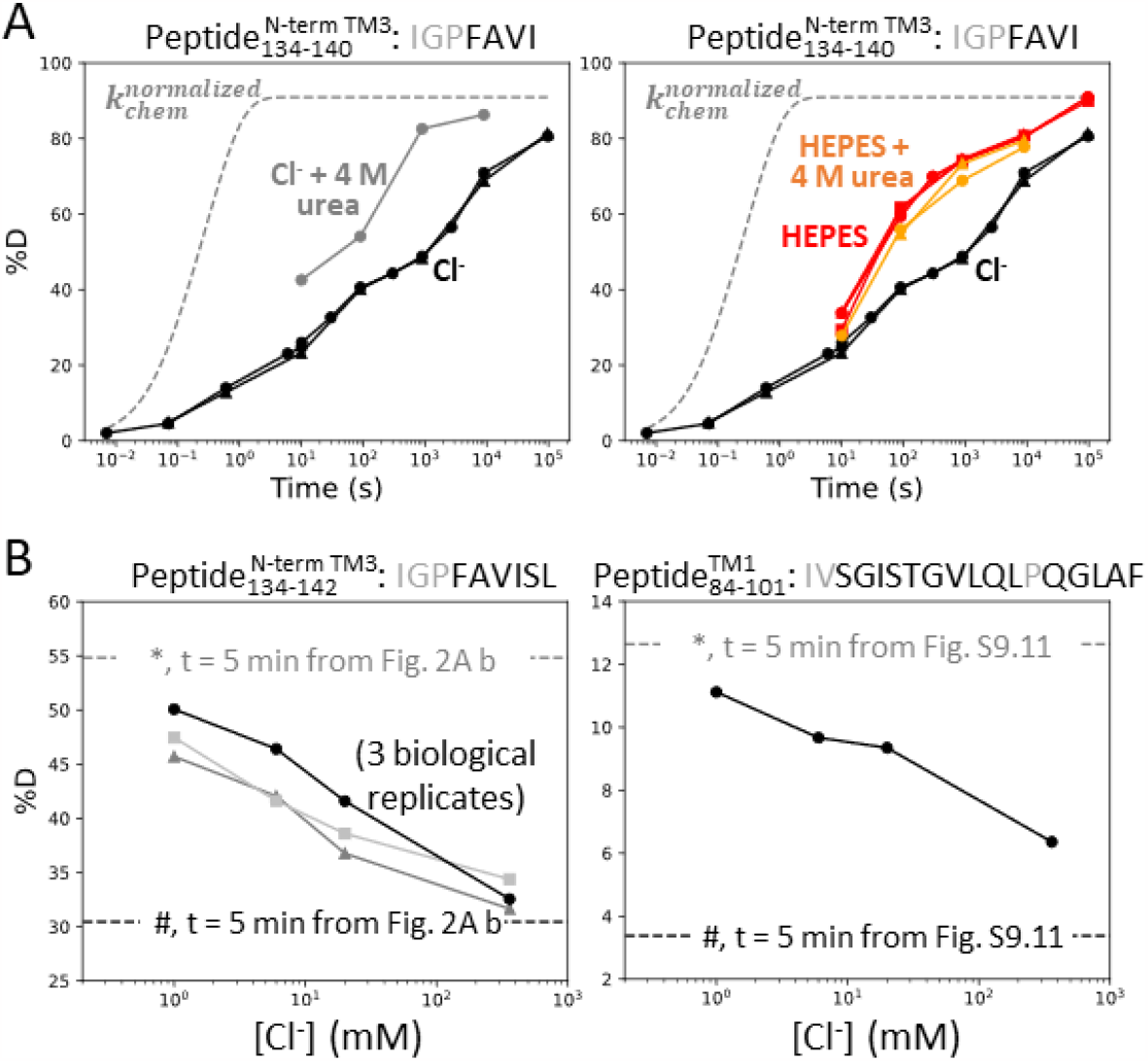
Anion binding folds and stabilizes prestin’s binding site. **(A)** Deuterium uptake plots for the N-terminus of TM3 (Peptide_134-140_) measured in the absence and presence of 4 M urea, in a background of (**Left**) Cl^-^ and (**Right**) HEPES. Replicates (circles, triangles, and squares): 2 in Cl^-^, 3 in HEPES, 2 in HEPES with urea, biological. Grey dashed curves represent deuterium uptake with *k*_*chem*_, normalized with the back-exchange level. **(B)** Deuteration levels for (**Left**) the N-terminus of TM3 (Peptide_134-142_) in three biological replicates and for (**Right**) TM1 (Peptide_84-101_) after 5 min labeling upon titrating Cl^-^ to apo state of prestin. Dashed lines indicate deuteration levels at t = 5 min (* and # for apo and Cl^-^-bound states, respectively) taken from **Fig. 2A.b** and **Fig. S9.11**. Residues in grey denoted in the peptide sequence do not contribute to the deuterium uptake curve.

We note that in apparent contradiction to our inference that the N-terminus of TM3 was unfolded in the apo state, its HDX was ∼100-fold slower than *k*_*chem*_. Such apparent PF for an unfolded region has been reported when it is located inside an outer membrane *β*-barrel, rationalized by the region having a lower effective local concentration of the HDX catalyst, [OD^-^], than in bulk solvent^25,26^. For prestin, we propose that detergent molecules in the micelle can restrict the access of OD^-^ to amide protons, leading to a local effective pD lower than the bulk solvent and hence producing the apparent PF for the unfolded N-terminus of TM3.

The folding reversibility of the anion-binding site was evaluated by tracking the HDX for 5 min after titrating in Cl^-^ to apo prestin. Deuteration levels for the N-terminus of TM3 (Region_137-140_) decreased with increasing Cl^-^ concentration (**Fig. 3B**), suggesting reversible folding upon Cl^-^ binding. Similar behavior was seen in the middle of TM1 (Region_86-101_: Peptide_84-101_) as Cl^-^ binding stabilized the binding pocket (**Fig. 3B**).

We also examined prestin’s stability in its intermediate states, obtained by replacing Cl^-^ anions with SO_4_^2−^ and salicylate^5,6^. When SO_4_^2−^ is the major anion, prestin’s HDX was nearly identical as in Cl^-^, except for a slightly faster HDX at the anion-binding site (Region_128-142_ and Region_394–397_) at labeling times longer than 10^3^ s (**Fig. 4**). This mild HDX response suggested a slightly destabilized binding site while the remaining regions retained normal dynamics as in Cl^-^. In the presence of salicylate, HDX slowed across the TMD, with the greatest effect seen at the anion-binding site (10-fold; 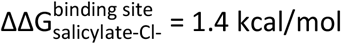) (**Fig. 4**), indicating that salicylate binding to prestin globally stabilized the TMD, primarily at the anion-binding site. These stability changes provide a thermodynamic context to the cryo-EM structures^5^.

**Figure 4:**
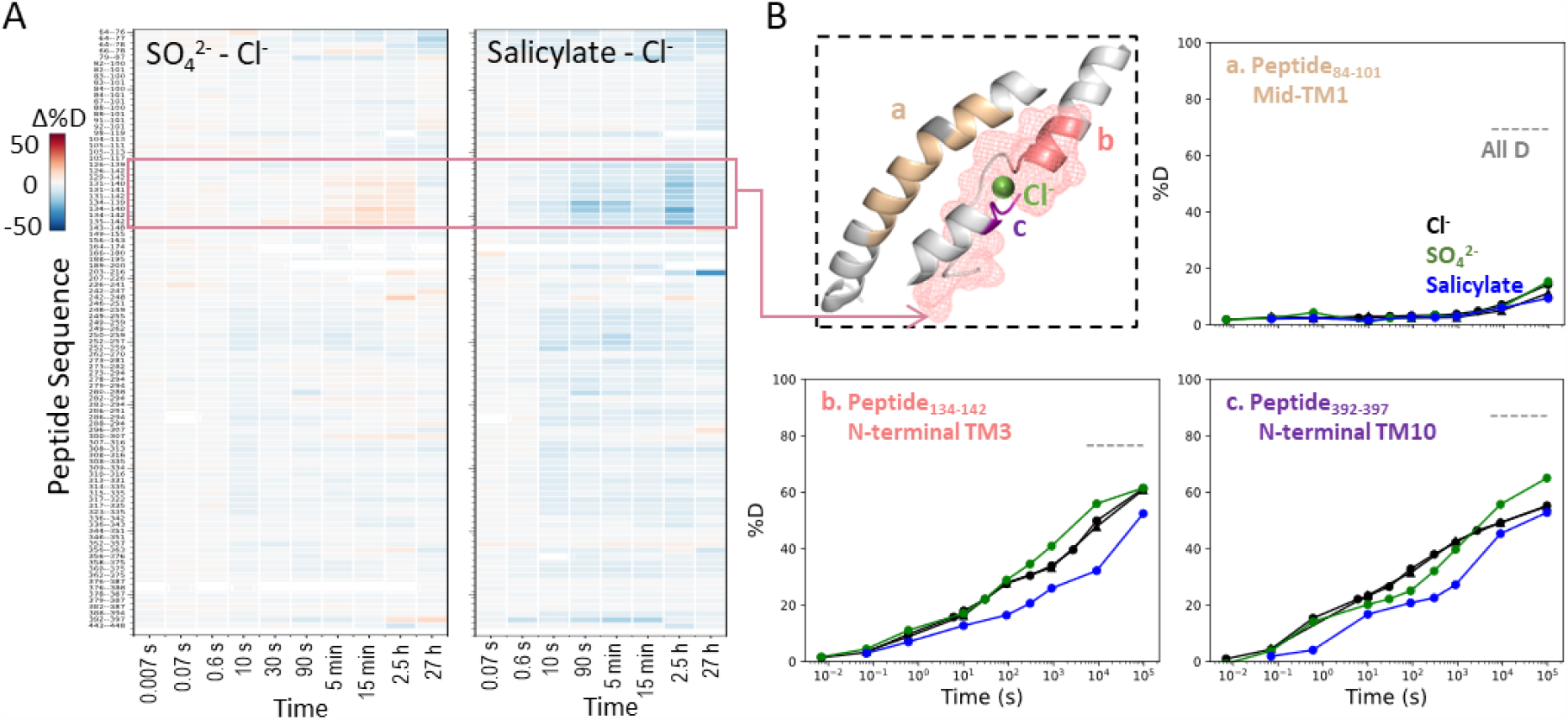
Prestin’s dynamics are regulated by anions of varying identities. **(A)** Heatmaps showing the difference in deuteration levels at each labeling time for all TMD peptides measured in SO ^2-^ or salicylate compared to Cl^-^. Peptide sequences are displayed on the y-scale and legible through the high resolution image. **(B)** The structure shows the anion-binding pocket with the putative position of the bound Cl^-^. The pink mesh highlights the region with the greatest HDX response to binding to various anions. Colored regions correspond to peptides whose deuterium uptake plots are shown when the protein is in Cl^-^ (black, two biological replicates shown in circles and triangles), SO ^2-^ (green), and salicylate (blue). Prolines are colored in grey. Grey dashed lines indicate deuteration levels in the full-D control.

### Prestin in a lipid bilayer exhibits a highly dynamic TM6

We chose to measure the HDX of prestin in GDN micelle to match the cryo-EM conditions^5–7^. Structures of GDN-solubilized prestin in Cl^-^ obtained by three research groups are indistinguishable (RMSD < 1 Å), demonstrating the reproducibility and robustness of the system.

To evaluate the dynamics in a more native membrane environment, we measured HDX of prestin reconstituted in nanodisc (porcine brain total lipid extract). Except for TM6, HDX for prestin in nanodisc highly resembled that in micelles including the anion-binding pocket (**Fig. 5**). Such high agreement between the folding stability in these two membrane mimetics suggest that our findings on prestin’s anion-binding site and its folding equilibrium are pertinent to prestin residing in a lipid bilayer. This is not surprising because structures for human prestin in GDN and nanodisc are shown to be nearly identical (Cα RMSD = 0.2 Å)^6^.

**Figure 5:**
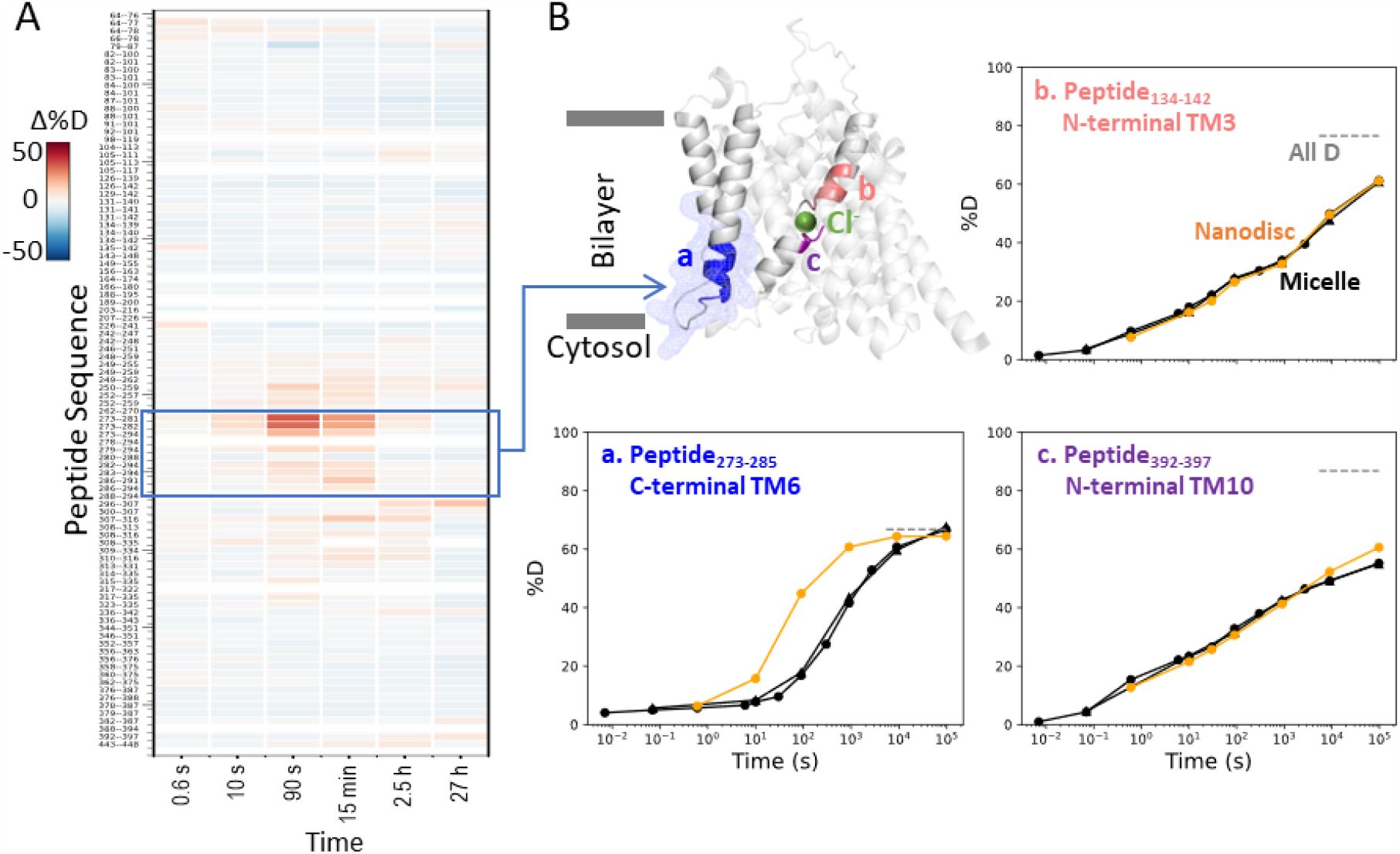
Prestin in nanodisc displays similar folding stability to prestin in micelle, except for a more dynamic TM6. **(A)** Heatmaps showing the difference in deuteration levels at each labeling time for all available TMD peptides measured for prestin in nanodisc (porcine brain total lipid extract) compared to prestin in detergent micelle (GDN), both in Cl^-^. Peptide sequences are displayed on the y-scale and legible through the high resolution image. **(B)** The structure shows the TMD for one subunit of prestin with the putative position of the bound Cl^-^. The blue mesh highlights the region where the greatest HDX difference was seen for prestin in nanodisc compared to micelle. Colored regions correspond to peptides whose deuterium uptake plots are shown when the protein is in micelle (black, biological duplicates shown in circles and triangles) and in nanodisc (orange). Grey dashed lines indicate deuteration levels in the full-D control.

Interestingly, nanodisc-embedded prestin displayed a less stable TM6 for the intracellular portion than prestin in micelles, manifested by the 10-fold HDX increase (Region_275-282_: Peptide_273-282_ + 2 peptides) (**Fig. 5 & Fig. S9.53-9.55**). TM6 defines the interface between prestin and the lipid bilayer, and has been proposed to mediate area expansion through helical bending^5^. The exact role of TM6 in regulating prestin’s conformational cycle is currently under investigation.

### Incremental unfolding of prestin’s binding site versus cooperative unfolding of the lipid-facing helices

Our broad HDX time range and dense sampling allowed us to observe effects at the residue level. In particular, the binding site of prestin (Region_128-140_ and Region_394–397_) exhibited a broad deuterium uptake curve in Cl^-^, indicative of helix fraying where exchange of deuterium occurs from multiple states that differ by one hydrogen bond (**Fig. 6A**). Such HDX pattern is consistent with the associated residues undergoing sequential unfolding with distinct PFs (i.e., stability). Site-resolved PFs obtained using PyHDX^21^ support that the stability increased residue-by-residue for TM3 for amide protons located further away from the substrate (**Fig. S3A**). This gradual increase in residue stability along the helices is indicative of helix fraying starting from prestin’s binding site.

**Figure 6:**
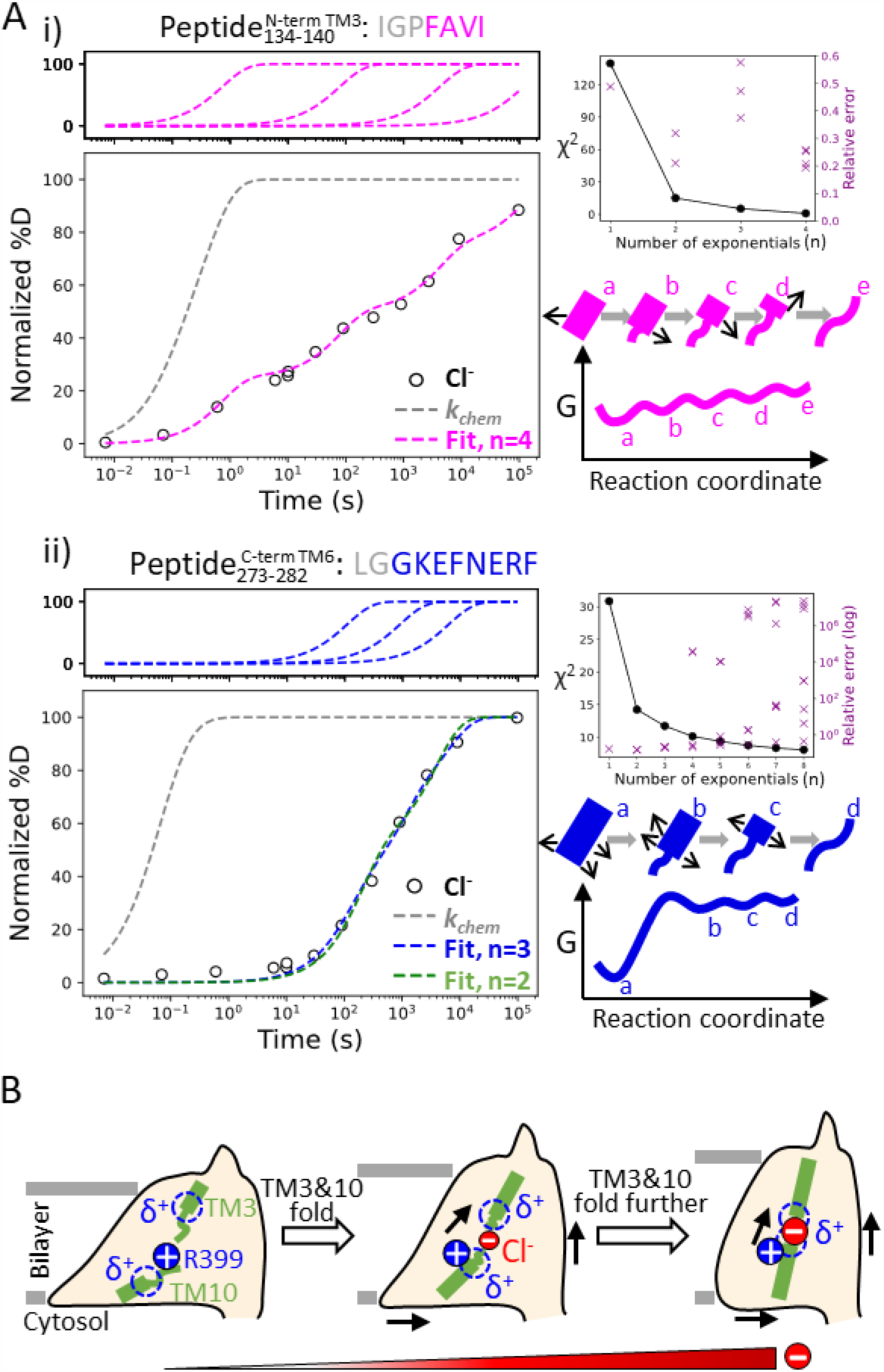
Helix folding cooperativity and the proposed mechanism for prestin’s electromotility. **(A) Left:** Deuterium buildup curves for **(i)** the N-terminal TM3 (Peptide_134-140_) and **(ii)** the intracellular portion of TM6 (Peptide_273-282_) in Cl^-^ depicting helix fraying and mild cooperativity, respectively. Circles: experimental deuteration levels, normalized with in- and back-exchange levels. Grey dashed curves: hypothetical intrinsic uptake curves (PF = 1). On the top shows individual exponentials whose sum is fitted to the experimental values and plotted on the main buildup curves. Residues in grey denoted in the peptide sequence do not contribute to the deuterium uptake curve. **Upper right:** χ^2^ and the relative error as the number of fit exponentials increases, used to assess the quality of fit. **Lower right:** Models and free energy surface of unfolding illustrating the difference between **(i)** fraying and **(ii)** mild cooperativity. Mechanism for prestin’s conformational transition from the expanded to the contracted state regulated by the anion concentration. Green rectangles and curved lines: folded and unfolded fractions, respectively, of TM3 and TM10. Blue filled circle: R399. Blue dashed circle: partial positive charges from TM3 and TM10 helical dipoles. Red filled circle: anions, with the size of the circle depicting anion concentrations. Black arrows: prestin’s conformational change.

In contrast, we observed much more cooperative unfolding in prestin’s lipid-facing helices, with exchange occurring from one or a few high energy states where a set of hydrogen bonds are broken concertedly. Cooperatively exchanging residues have similar PFs and a characteristic sigmoidal deuterium uptake curve for the associated peptide, as seen in prestin’s intracellular portion of TM6 (Region_275-282_: Peptide_273-282_ + 4 peptides) (**Fig. 6A & Fig. S9.53-9.57**).

To characterize the degree of cooperativity for the HDX at the N-terminus of TM3 (Peptide_134-140_) and the intracellular portion of TM6 (Peptide_273-282_), we fit the deuterium uptake curves as a sum of exponentials^17^, 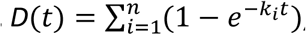, where *k*_*i*_ is the exchange rate and *n* is the number of exponentials, ranging from one to the number of exchange-competent residues. The value of *n* was determined by the quality of the fit, evaluated by χ^2^ and having a relative error smaller than one (i.e., standard deviation for *k*_*i*_less than *k*_*i*_itself). HDX data for the N-terminus of prestin’s TM3 (Peptide_134-140_) was fit with four well-separated exponentials with rates spanning five log units for the four residues (**Fig. 6A**). The need to individually fit each site indicates a lack of cooperativity and helix fraying. In contrast, the peptide representing the intracellular portion of TM6 (Peptide_273-282_) has eight residues yet it could be fitted with only three rates spanning less than two log units (**Fig. 6A**). This rather concerted deuterium uptake was independent of the anion substrate identity and also observed for TM1, TM5b, the intracellular portion of TM7, and the N-terminus of TM8 (**Fig. S9.6-9.16, S9.42-9.43, S9.45-9.51, S9.58-9.63, S9.78-9.79**).

We define a parameter *σ*_*ΔG*_ to quantify the degree of folding cooperativity. The value of *σ*_*ΔG*_ is calculated as the standard deviation for the free energies of unfolding (ΔGs) for exchange-competent residues comprising the peptide. When a region folds 100% cooperatively, *σ*_*ΔG*_ is zero as all residues have the same ∆G. As the diversity increases (lower cooperativity), the *σ*_*ΔG*_ value becomes larger. The accuracy of the ∆G determination at residue level can be increased by comparing uptake curves for overlapping peptides and/or deconvoluting isotope envelopes^17^. Here we assigned exchange rates (*k*_*i*_), obtained from the fitting method mentioned above, to residues based on the directionality of helix fraying, with residues closer to the end of a helix having faster rates. When there is ambiguity on which rate to assign to a given residue, the geometric mean of the rates was used (**Materials and Methods**). We found that prestin’s intracellular portion of TM6 (Peptide_273-282_) has *σ*_*ΔG*_ = 1.1, indicating mild cooperativity, whereas the non-cooperative N-terminal TM3 (Peptide_134-140_) has a *σ*_*ΔG*_ = 2.9. This significant decrease in folding cooperativity for helices directly involved in the Cl^-^ binding site likely has functional consequences related to prestin’s electromotility, as discussed below.

## Discussion

Using HDX-MS, we provide novel information on the structural dynamics of prestin in its apo state, for which there isn’t an associated cryo-EM structure. We demonstrate that prestin displays very similar dynamics in nanodisc as in micelles, except for a destabilized lipid-facing helix TM6 that is critical for mechanical expansion. We have explored the energetic and conformational differences between prestin, a voltage-dependent motor, and its mammlian relative SLC26A9, a representative member of the SLC26 family of anion transporters for which a cryo-EM structure is available. Our data point to major differences in the energetics at the anion-binding site of prestin and SLC26A9 despite their structural similarities. This comparison addresses underlying mechanistic questions related to the unique properties of prestin, the origin of its voltage dependence, and the potential mechanisms that couple charge movements to electromotility.

We showed that prestin displays an unstable binding site, regardless of being in nanodisc or micelles (**Fig. 5**). Upon Cl^-^ unbinding, the binding site unfolds by one helical turn at the electrostatic gap formed by the abutting (antiparallel) short helices TM3 and TM10 (**Fig. 2A & Fig. 3**). We measured an increase in local ∆∆G = 1.8∼3.5 kcal/mol upon anion binding. This energy difference is within the range of the ∆∆G = 2.4 kcal/mol estimated from having a 60-fold excess of Cl^-^ above the EC_50_ (6 mM)^11^. Similar folding events upon anion binding are absent in SLC26A9 (**Fig. 2B**), pointing to a key role of the bound anion as a structural element in prestin, stabilizing the natural repulsion between TM3-TM10 positive helical macrodipoles. This phenomenon rationalizes the conundrum that prestin’s voltage dependence requires the proximity of a bound anion to R399.

We find that anion binding to prestin mainly stabilizes the interface between the scaffold and the elevator domains (**Fig. 1C & Fig. S1A**). This phenomenon is consistent with an elevator-like mechanism during prestin’s conformational transition from the expanded to the contracted state^5^. Anion binding stabilizes prestin’s intracellular cavity and slightly destabilizes regions facing the extracellular matrix. This effect can result from changes in solvent exposure, as the intracellular water cavity may shrink as prestin contracts. For SLC26A9, the destabilization upon anion binding at the intracellular cavity likely results from a shift from the outward-facing state to the inward-facing state (**Fig. 1C & Fig. S1B**), supporting the alternate-access mechanism for this fast transporter^9,10^. Similar HDX changes, i.e., increased HDX on the intracellular side while decreased HDX on the extracellular side, have been observed in other transporters during their transition from outward-facing to inward-facing states^27,28^. Prestin’s distinct HDX response compared to a canonical transporter is consistent with it being an incomplete anion transporter^11,29^.

The HDX data for prestin in Cl^-^, SO_4_^2−^, and salicylate support an allosteric role for the anion binding at the TM3-TM10 electrostatic gap^12–14^ (**Fig. 4**). SO_4_^2−^ binding leads to shifts in the NLC towards positive potentials, thus stabilizing multiple conformations that are on average more expanded than prestin in Cl^-5,13,30^. Since the binding of SO_4_^2−^ to prestin is weaker than that of Cl^-6^, the slight increase in HDX at the binding site likely reflects more prestin molecules adopting the apo state. Salicylate binding inhibits prestin’s NLC and yet the molecular basis of such inhibition remains obscure^5,16^. Bavi *et al*^5^ showed that binding of salicylate occludes prestin’s binding pocket from solvent and inhibits the movement of TM3 and TM10. This is fully consistent with the 10-fold HDX slowing found for the N-termini of TM3 and TM10 upon salicylate binding as compared to the rest of the protein. Our HDX data, together with results from Bavi *et al*.^5^, suggest that salicylate likely inhibits prestin’s NLC by restricting the dynamics of the anion-binding site.

We identified helix fraying at the anion-binding site of prestin based on its broad deuterium uptake curve in the presence of Cl^-^, consistent with a multi-state landscape (**Fig. 6A**). This fraying suggests that an increase in the anion concentration would promote helical propensity at TM3 and TM10, and is inconsistent with a cooperative (two-state) model involving an equilibrium between an apo state and a single bound state (**Fig. 6A**). Therefore, we suggest that a two-state model of prestin’s conformational changes with a high energy barrier would be insufficient to explain its fast kinetics, whereas charge movement is facilitated by crossing multiple shallow barriers^13,31^.

We propose that having a Cl^-^ binding site that frays can promote prestin’s fast motor response which is thought to have evolved independently of its voltage sensing ability^32,33^. While the stability for TM10 is similar for prestin and SLC26A9, the latter protein exhibits a more stable, non-fraying TM3 (**Fig. 2**). Notably, the normally highly conserved Pro136 in mammalian prestin is replaced with a Threonine in SLC26A9 and other vertebrates that express non-electromotile prestin (**Fig. S4A**). This Pro136Thr substitution, based on our *Upside* MD simulations, results in a hyper-stabilized TM3 that would otherwise have similar folding stability as TM10 (**Fig. S4B**). A Pro136Thr mutation in rat prestin also leads to a shift of NLC towards depolarized potentials^20^. These thermodynamic and functional consequences of having a destabilized TM3 with Pro136, which now has similar stability as TM10, lead us to hypothesize that prestin’s fast mechanical activity may be promoted by having simultaneous fraying of the TM3 and TM10 helices.

Helices that exhibit cooperative unfolding all appear to be lipid-facing helices, including TM6-TM7, TM1, TM5b, and TM8. The region with the most pronounced cooperativity, the intracellular portion of TM6, has a series of glycines including the consecutive G274-G275 pair that underlies the “electromotility elbow”, a helical bending contributing to the largest cross-sectional area (expanded conformation) and the thin notch in the micelle^5^. Importantly, HDX for prestin in nanodisc reveals a significant stability decrease at TM6 while the remaining regions retained similar dynamics as in micelles. This high sensitivity of TM6 stability to membrane environment, together with the structural consequences of cooperativity, speak to the its significance in prestin’s area expansion. We propose that cooperativity allows for long-range allostery^34^ so that the lipid-facing helices, particularly TM6, can adopt large-scale structural rearrangements as induced by voltage sensor movements, thereby achieving rapid electromechanical conversions of prestin. The exact mechanism through which cooperativity contributes to prestin’s electromotility remains a key question.

Based on the structural and allosteric role of Cl^-^ binding at the TM3-TM10 electrostatic gap, we propose a model in which prestin’s conformational changes and electromotility are regulated by the folding equilibrium of the anion-binding site (**Fig. 6B**). In our model, anion binding participates in a local electrostatic balance that includes the positively charged R399 and the positive TM3-TM10 helical macrodipoles. In the apo state, the anion-binding site unfolds due to the electrostatic repulsion between these positively charged groups. Being a buried charge, R399 may exit from the electric field concentrated in the lower dielectric environment of the protein, and move into the solvent region as it lacks a neutralizing anion. This event is coupled to the allosteric expansion of prestin’s membrane footprint^5^. Anion binding partially neutralizes the positive electric field at the binding site, an event that is coupled to the residue-by-residue folding for the N-termini of TM3 and TM10 as well as the shortening of the electrostatic gap. This folding event results in a more focused electric field and consequent contraction of prestin’s intermembrane cross-sectional area. At physiologically-relevant low Cl^-^ concentration (1.5-4 mM)^35^, prestin’s binding site is likely to be only partially folded. Complete folding may be achieved by membrane potential acting on the TM3-TM10 helical dipoles, which leads to the movement of the voltage sensor across the electric field and the rapid areal expansion for the TMDs (i.e., electromotility).

### Structure of prestin in HEPES and low Cl^-^ levels

We associate the HEPES solvent condition to an apo state as the HEPES anion is too big to fit into the chloride site and prestin has a right-shifted NLC^13^, To investigate further whether HEPES anion binds, we determined the structure of prestin in the HEPES-based buffer using cryo-EM. Our initial attempt to solve the structure in the complete absence of Cl^-^ was unsuccessful due to sample aggregation under cryogenic conditions. Aggregated particles were greatly reduced in the presence of 1 mM Cl^-^, and we were able to solve the structure of GDN-solubilized prestin in 1 mM Cl^-^, containing 190 mM HEPES, at a nominal resolution of 3.4 Å (**Supporting Information Text 3, Fig. S11 & Fig. S12**). Surprisingly, under these conditions prestin adopted a “compact” conformation, virtually identical to the previously reported Cl^-^-bound “Up” state^5^, displaying a clear density at the anion binding site (**Fig. S11C**). This density is incompatible with the placing of a HEPES molecule, and we reason it to be a small population of Cl^-^-bound prestin resulting from a weak Cl^-^ affinity (e.g., EC_50_=6 mM^11^ implies 17% bound). This result is consistent with the fundamental role of bound anions in the conformational stability of prestin and supports a new role for the folding equilibrium of the anion-binding site in the mechanism of voltage sensing. Ultimately, however, understanding the underlying mechanism for prestin’s electromotility necessitates consideration of physiological elements such as membrane potential, kHz frequency, and protein-lipid interactions.

## Conclusions

We applied HDX-MS to the study of prestin’s electromotility and identified folding events that are likely critical for function but had escaped detection by cryo-EM. The folding equilibrium of the Cl^-^ binding site and its dependence on Cl^-^ concentration appears to rationalize the conundrum of how an anion that binds in proximity to a positive charge (R399), can enable the NLC of prestin. We directly compared the dynamics of prestin in nanodisc to in micelles and identified TM6 as a potential mechano-sensing helix. We observed fraying of the helices forming the anion-binding site, which contrasts with cooperative unfolding of the lipid-facing helices. We believe that the non-cooperative fraying of the helices involved in voltage sensing may allow for fast charge movements within the electric field. This heightened sensitivity of the voltage sensor then induces large-scale motions of the lipid-facing helices, enabled by their cooperativity (or allostery), thereby altering the cross-sectional area of prestin. These principles warrant further investigation.

## Materials and Methods

### Sample preparation for prestin and SLC26A9

Generation of the dolphin prestin and mouse SLC26A9 constructs, protein overexpression, and purification were conducted using the same protocol as previously described^5^. Following the cleavage of the GFP tag, the protein was purified by size-exclusion chromatography (SEC) on a Superose 6, 10/300 GE column (GE Healthcare), with the running buffer being either the “Cl^-^/H_2_O Buffer” or the “SO_4_^2−^/H_2_O Buffer”, including 1 μg/mL aprotinin and 1 μg/mL pepstatin^5^. The “Cl^-^/H_2_O Buffer” contained 360 mM NaCl, 20 mM Tris, 3 mM dithiothreitol, 1 mM EDTA, and 0.02% GDN at pH 7.5. The “SO_4_^2−^/H_2_O Buffer” contained 140 mM Na_2_SO_4_, 5 mM MgSO_4_, 20 mM Tris, 0.02% GDN, and 10-15 mM methanesulfonic acid to adjust the pH to 7.5. Peak fractions containing the sample were concentrated on a 100K MWCO centrifugal filter (Millipore) to 2-3 mg/mL, flash-frozen in liquid nitrogen, and kept at *-*80 °C until use.

For prestin reconstitution into nanodiscs, porcine brain lipid extract (Avanti) was first solubilized in “Cl^-^/H2O Buffer” supplemented with 3% GDN to a 10 mg/mL concentration through successive rounds of sonication and freeze-thaw cycles. MSP1E3D1 was purified as previously described^36^.^37^ Reconstitution was performed using the “on-column” method^37^ using a 1:5:600 (prestin:MSP:lipid) molar ratio, with the final buffer consisting of 20 mM Tris-HCl and 150 mM NaCl at pH 7.5.

For structure determination by cryo-EM, prestin purification was performed in buffers containing 360 mM NaCl as previously described^5^, except that the SEC running buffer now consisted of 190 mM HEPES, ∼95 mM Tris-base (used to adjust the pH to 7.5), 1 mM NaCl, 3 mM DTT, 0.02% GDN, 1 μg/mL aprotinin, and 1 μg/mL pepstatin. Peak fractions containing the sample were concentrated to 3 mg/ml and used immediately for grid preparation.

### Hydrogen-deuterium exchange

**Table S1** provides biochemical and statistical details for HDX in this study per recommendations by Masson *et al*^38^. HDX reactions, quench, and injection were all performed manually. Prior to HDX, proteins purified in Cl^-^ and SO_4_^2−^ were buffer exchanged to the same H_2_O buffer without protease inhibitor using 7K MWCO Zeba spin desalting columns (Thermo 89882). For HDX conducted in HEPES, proteins purified in Cl^-^ were dialyzed against the “HEPES/H_2_O Buffer” (150 mM HEPES, 0.02% GDN, pH adjusted to 7.5 by HEPES acid or base) using a 10K MWCO dialysis device (Thermo Slide-A-Lyzer MINI 69570) with three times of buffer exchange for near-complete Cl^-^ removal. HDX in a solution of 93% deuterium (D) content was initiated by diluting 2 μL of 25 μM prestin or SLC26A9 stock in an H_2_O buffer into 28 μL of the corresponding buffer made with D_2_O (99.9% D, Sigma-Aldrich 151882). HDX was conducted in one of the two conditions: 1) pD_read_ 7.1, 25 °C; 2) pD_read_ 6.1, 0 °C. The D_2_O buffers contained the same compositions as the corresponding H_2_O buffers, except that Tris was replaced with Phosphate for the “SO_4_^2−^/D_2_O Buffer” for both HDX conditions, and the “Cl^-^/D_2_O Buffer” for HDX performed at pD_read_ 6.1, 0 °C. The pD_read_ was adjusted to 7.1 or 6.1 by DCl for the “Cl^-^/D_2_O Buffers” and by NaOD for other D_2_O buffers. For HDX in the presence of salicylate, 50 mM salicylate acid was added to the “SO_4_^2−^/D_2_O Buffer” and the pD_read_ was adjusted by NaOD. For HDX in the presence of urea, 4 M urea was added to the “Cl^-^/D_2_O Buffer” or the “HEPES/D_2_O Buffer”, with accurate urea concentration determined to be 4.16 M and 4.54 M, respectively, by refractive index using a refractometer (WAY Abbe)^39^.

HDX was quenched at various times, ranging from 1 s to 27 h, by the addition of 30 μL of ice-chilled quench buffer containing 600 mM Glycine, 8 M urea, pH 2.5. For HDX in the presence of urea, urea concentration in the quench buffer was adjusted to reach a 4 M final concentration. For HDX on prestin in nanodisc, the quench buffer also included 3 μL of 0.8% GDN and 3 μL of 300 mg/ml aqueous suspension of ZrO2-coated silica (Sigma-Aldrich, reference no. 55261-U). The resulting mixture was incubated on ice for 1 min to remove lipids and solubilize prestin in GDN before being filtered through a cellulose acetate spin cup (Thermo Pierce, Waltham, Massachusetts, reference no. 69702) by centrifugation for 30 s at 13,000 g, 2 °C. Quenched reactions were immediately injected into a valve system maintained at 5 °C (Trajan LEAP). Non-deuterated controls and MS/MS runs for peptide assignment were performed with the same protocol as above except D_2_O buffers were replaced by H_2_O buffers, followed by the immediate addition of the quench buffer and injection.

HDX reactions were performed in random order. No peptide carryover was observed as accessed by following sample runs with injections of quench buffer containing 4 M urea and 0.01% GDN. In-exchange controls accounting for forward deuteration towards 41.5% D in the quenched reaction were performed by mixing D_2_O buffer and ice-chilled quench buffer prior to the addition of the protein. Maximally labeled controls (“All D”) accounting for back-exchange were performed by a 48-h incubation with the “Cl^-^/D_2_O Buffer” at 37 °C, followed by a 30-min incubation with 8 M of deuterated urea at 25 °C.

### Protease digestion and LC-MS

Upon injection, the protein was digested online by a pepsin/FPXIII (Sigma-Aldrich P6887/P2143) mixed protease column maintained at 20 °C. Protease columns were prepared in-house by coupling the protease to a resin (Thermo Scientific POROS 20 Al aldehyde activated resin 1602906) and hand-packing into a column (2 mm ID × 2 cm, IDEX C-130B). After digestion, peptides were desalted by flowing across a hand-packed trap column (Thermo Scientific POROS R2 reversed-phase resin 1112906, 1 mm ID × 2 cm, IDEX C-128) at 5 °C. The total time for digestion and desalting was 2.5 min at 100 μL/min of 0.1% formic acid at pH 2.5. Peptides were then separated on a C18 analytical column (TARGA, Higgins Analytical, TS-05M5-C183, 50 x 0.5 mm, 3 μm particle size) via a 14-min, 10–60% (vol/vol) acetonitrile (0.1% formic acid) gradient applied by a Dionex UltiMate-3000 pump. Eluted peptides were analyzed by a Thermo Q Exactive mass spectrometer. MS data collection, peptide assignments by SearchGUI version 4.0.25, and HDX data processing by HDExaminer 3.1 (Sierra Analytics) were performed as previously described^25,26^.

### HDX data presentation, quantification, and statistics

In our labeling convention, we name capitalized regions according to the third residue of each peptide since the first two residues have much faster *k*_chem_ ^18,19^ and hence, exhibit complete back-exchange. Labeling times for HDX performed in pD_read_ 6.1, 0 °C were corrected to those in pD_read_ 7.1, 25 °C by a factor of 140, determined by the ratio of the *k*_*chem*_ for full-length proteins in the two conditions – prestin: 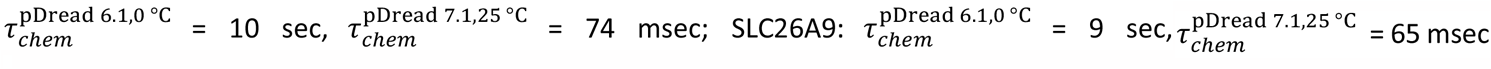. Deuteration levels were adjusted with the 93% D content but not with back-exchange levels except for **Fig. 6A**. For HDX in the presence of urea (**Fig. 3A**), D contents of 76% and 74% were used to account for the volumes of 4.16 M and 4.54 M urea in the “Cl^-^/D_2_O Buffer” and the “HEPES/D_2_O Buffer”, respectively.

HDX rates, protection factors (PFs), and changes of free energy of unfolding (ΔΔG’s) in the Results section were estimated by fitting uptake curves of each peptide, after correcting for back-exchange levels, with a stretched exponential as described by Hamuro 2021^17^, except for discussion relevant to **Fig. 6A**. Peptides with less than 10% D at the longest labeling time (∼10^5^ s) were not used for fitting. The residue-level ΔΔG values presented in the full-length proteins in **Fig. 1C** were estimated by averaging ΔΔG values for peptides containing the residue. We note that this stretched exponential method is only a rough approximation to extract ΔΔG’s and our major conclusions are not dependent on this fitting method.

For fitting HDX data and extracting rates relevant to **Fig. 6A**, HDX data were first normalized to in- and back-exchange levels. Given the helices are likely to exchange by fraying, the exchange rate for each residue was assigned based on their distance from the end of the helix. When a residue could not be assigned to a single rate, the geometric mean of the possible rates was used, e.g., the HDX data for the 8-residue Peptide_273-282_ were well fit with 3 exponentials, and the three associated rates, k_1_, k_2_, & k_3_, were assigned to the 8 residues according to k_1_, k_1_, (k_1_k_1_k_2_)^1/3^, k_2_, k_2_, (k_2_k_2_k_3_)^1/3^, k_3_, & k_3_. These rates were used to calculate folding stability according to △G = *-*RTln(*k*_*Chem*_/*k*_*i*_*-* 1).

A hybrid statistical analysis used to generate **Fig. S1** was performed as described by Hageman & Weis, 2019^40^, with significance limits defined at α = 0.05.

### MD simulations

Simulations were conducted in our Upside molecular dynamics package^41,42^ using a membrane thickness of 38 Å. Missing residues for prestin (PDB 7S8X) were built using MODELLER^43^ and the placement within the bilayers was accomplished using Positioning of Proteins in Membranes webserver^44^. Local restraints in the form of small springs between nearby residues were used to maintain the native structure of cytosolic domains, as well as to the TM13-TM14 helices. Also, the distance between the two TMDs was held fixed. We ran 28 temperature replicas between 318 and 360K.

### Cryo-EM sample preparation and data collection

The cryo-EM data was collected from three separate samples and microscope sessions. For dataset 1, 3.5 μL of prestin sample was applied to Quantifoil 200-mesh 1.2/1.3 Cu grids (Quantifoil) that were plasma cleaned for 30s at 20W. For datasets 2 and 3, 3.5 μL of prestin sample was applied to UltrAuFoil 300-mesh 1.2/1.3 grids (Quantifoil UltrAuFoil) that were plasma cleaned for 40s at 20W. The remaining sample preparation and imaging conditions were kept constant for all three samples. Grids were blotted at 22 °C and 100% humidity with a blot time of 3.5 s and a blot force of 1, and flash-frozen into liquid ethane using a Vitrobot Mark IV (Thermo Fisher). Grids were imaged at The University of Chicago Advanced Electron Microscopy Facility on a 300 kV Titan Krios G3i electron microscope equipped with a Gatan K3 camera in CDS mode, a GIF energy filter (set to 20 eV) and with magnification set to 81,000x, corresponding to a physical pixel size of 1.068 Å. Movies were acquired at a dose of 1.2 e^-^/Å^2^ for 50 frames (corresponding to a total dose of 60 e^-^/Å^2^) and a defocus range of -0.7 to -2.1 μm. 2,153 movies were collected for dataset 1, 1,928 movies for dataset 2, and 6,665 movies for dataset 3.

### Cryo-EM Image Processing

Cryo-EM data processing was performed using Relion-4.0^45^ and cryoSPARC v4.1^46^. Movies from the different datasets were motion-corrected independently in Relion with a bin-1 pixel size of 1.068 Å. The motion-corrected micrographs were then combined and imported into cryoSPARC for Patch CTF Estimation. Unless otherwise mentioned, the following steps were performed in cryoSPARC. Particles were picked using template-based particle picking. The initial picks were curated using Inspect Picks and extracted at a box size of 256. Two rounds of 2D Classification were performed to filter out additional “junk” particles, resulting in a set of 328,033 particles. Two consecutive rounds of ab initio reconstruction, followed by heterogeneous refinement with C1 symmetry were performed to obtain a stack of 170,313 particles. These particles were used as input for an initial round of non-uniform and local CTF refinement using C2 symmetry and exported to Relion for Bayesian Polishing. Additional 3D Classification was performed in Relion to remove additional “junk” particles, resulting in a set of 170,313 particles. However, no additional classes were found. The final set of polished particles was then imported back into cryoSPARC and subjected to a final round of non-uniform and CTF refinement, resulting in a map with a nominal resolution of 3.4 Å, according to the gold-standard 0.143 FSC criterion^47^. Local resolution was estimated using local resolution estimation in cryoSPARC.

### Model building and refinement

A previous model of dolphin prestin (PDB 7S8X) was roughly fit into the density map and used as a template for model building. Initially, only a monomer was considered for model building, and the fitting was improved by running Phenix real space refinement^48^ with secondary structure restraints, morphing, and simulated annealing enabled. Subsequently, the monomer model was iteratively refined by manual inspection in Coot^49^ and real space refinement without morphing and simulated annealing in Phenix. After several rounds of refinement, the second monomer was added using the apply_ncs tool in Phenix with C2 symmetry, and the resulting dimer model was subjected to an additional round of manual inspection in Coot. A chloride ion was manually added into the density observed in the anion binding pocket in Coot. All figures related to the cryo-EM structure were prepared using UCSF ChimeraX^50^.

## Description of supplementary material and file names

1. Supporting Information Text 1: Heterogeneity and HDX kinetics.
2. Supporting Information Text 2: Combining HDX-MS and cryo-EM in structural biology.
3. Supporting Information Text 3: Structure of prestin in HEPES and low Cl^-^ levels
4. Figure S1: Volcano plot analysis of HDX for prestin and SLC26A9 in response to Cl^-^ binding.
5. Figure S2: Site-resolved protection factors for prestin and SLC26A9 obtained using PyHDX.
6. Figure S3: PyHDX fitting supports that prestin exhibits helix fraying at the N-terminus of TM3 and mild cooperativity at the intracellular portion of TM6.
7. Figure S4: Mammalian prestin has a conserved and helix-destabilizing proline 136 on TM3.
8. Figure S5: HDX-MS sequence coverage and measurements for prestin and SLC26A9 in Cl^-^.
9. Figure S6: Regions unresolved in cryo-EM structures are unfolded in all conditions examined.
10. Figure S7: HDX for prestin occurs via EX2 mechanism.
11. Figure S8: Heterogeneity and HDX kinetics in TM1 and TM9.
12. Figure S9: Deuterium uptake curves for all peptides covering prestin’s transmembrane domain.
13. Figure S10: Deuterium uptake curves for all peptides covering SLC26A9’s transmembrane domain.
14. Figure S11: The cryo-EM structure for prestin in a HEPES-based buffer containing 1 mM Cl^-^ highly resembles the structure in the reported Cl^-^-bound state.
15. Figure S12: Workflow for the processing of the cryo-EM data.
16. Table S1: Biochemical and statistical details for HDX.

## Acknowledgments

This work was supported by grants to T.R.S. from the NSF (MCB 2023077) and the NIH (GM55694, 1R35GM148233), and to E.P. from R01 DC019833. We would like to thank the Advanced Electron Microscopy Facility at The University of Chicago for providing cryo-EM training and help with data collection.

## Data availability

The atomic structure coordinates have been deposited at the RCSB PDB under accession numbers 8UC1; and the EM maps have been deposited in the Electron Microscopy Data Bank under accession numbers EMD-42112. All materials generated during the current study are available from the corresponding author under a materials transfer agreement with The University of Chicago.

