## Supplementary Information for "Folding of Prestin’s Anion-Binding Site and the Mechanism of Outer Hair Cell Electromotility"

**Description of supplementary material and file names:**

1. Supporting Information Text 1: Heterogeneity and HDX kinetics.
2. Supporting Information Text 2: Combining HDX-MS and cryo-EM in structural biology.
3. Supporting Information Text 3: Structure of prestin in HEPES and low $Cl^-$ levels.
4. Figure S1: Volcano plot analysis of HDX for prestin and SLC26A9 in response to $Cl^-$ binding.
5. Figure S2: Site-resolved protection factors for prestin and SLC26A9 obtained using PyHDX.
6. Figure S3: PyHDX fitting supports that prestin exhibits helix fraying at the N-terminus of TM3 and mild cooperativity at the intracellular portion of TM6.
7. Figure S4: Mammalian prestin has a conserved and helix-destabilizing proline 136 on TM3.
8. Figure S5: HDX-MS sequence coverage and measurements for prestin and SLC26A9 in $Cl^-$.
9. Figure S6: Regions unresolved in cryo-EM structures are unfolded in all conditions examined.
10. Figure S7: HDX for prestin occurs via EX2 mechanism.
11. Figure S8: Heterogeneity and HDX kinetics in TM1 and TM9.
12. Figure S9: Deuterium uptake curves for all peptides covering prestin's transmembrane domain.
13. Figure S10: Deuterium uptake curves for all peptides covering SLC26A9's transmembrane domain.
14. Figure S11: The cryo-EM structure for prestin in a HEPES-based buffer containing 1 mM Cl- highly resembles the structure in the reported Cl--bound state.
15. Figure S12: Workflow for the processing of the cryo-EM data.
16. Table S1: Biochemical and statistical details for HDX

**Supporting Information Text 1: Heterogeneity and HDX kinetics.**

HDX in our study occurred mostly via EX2 kinetics where the observed exchange rate ($k_{ex}$) reports on the equilibrium (i.e., stability) rather than the opening rates of the exchange-competent states. The identification of EX2 behavior is supported by 1) $k_{ex}$ of prestin in the two HDX conditions differed by 140-fold, which can be attributed solely to the effect of pH and temperature on $k_{chem}$ for residues across the entire protein (**Fig. S7A**); 2) the continuous shifts in the single isotopic envelopes towards higher m/z with exchange time (**Fig. S7B**); 3) the envelopes had the binomial distribution expected when each site exchanges independently, supported by HDExaminer (v3.3) fits. The tracking of $k_{ex}$ with $k_{chem}$ also argues that prestin exhibits similar dynamics under the two HDX conditions, allowing us to combine the experiments after correcting for the difference in $k_{chem}$.

We observed bimodal isotopic envelopes for peptides in TM1 ($Region_{84-101}$: 9 peptides), with both envelopes increasing in mass over time, one exchanging slower than the other (**Fig. S8A**). Bimodality can result from HDX occurring via EX1 kinetics where every opening event results in exchange; this occurs when the rate of reforming the hydrogen bond, $k_{close}$, is much slower than $k_{chem}$. The signature of EX1 kinetics is a decrease in the amplitude of the lighter envelope and a commensurate increase in the heavier amplitude over time[1]. This EX1 behavior is observed in TM9 ($Region_{378-387}$) (**Fig. S8B**). Alternatively, bimodality can reflect the presence of two non- or slowly-interconverting, structurally distinct populations, each having its own exchange behavior. Peptides in TM1 retained a 1:3 ratio of relative intensity for the heavy-to-light envelopes regardless of biological replicates or anionic conditions, pointing to kinetically distinct populations. In addition, both populations in TM1 exchanged via EX2 kinetics (**Fig. S8A**). Therefore, conformational heterogeneity best explains the exchange behavior of TM1 with an interconversion time between the two populations being longer than our longest labeling time (27 h). We associate the slow population with the natively folded TM1, as observed in cryo-EM studies[2], and focus on this population in the present study.

As just noted, the presenting data for TM1 point to conformational heterogeneity with two populations having distinct HDX behavior. We believe that the fast population has TM1 unfolded despite it having a PF of ~100. We attribute this residual protection to detergent molecules hindering solvent access to the backbone and hence slowing exchange. This is the same explanation we provided for the heightened protection observed for the N-terminal TM3 ($Region_{137-140}$) in apo prestin (**Fig. 3A**). Generally, intrinsic HDX rates for unfolded regions of a soluble protein (i.e., $k_{chem}$) may not always serve as an appropriate reference rate for membrane-associated regions in the presence of detergents or lipids.

**Supporting Information Text 2: Combining HDX-MS and cryo-EM in structural biology.**

HDX-MS can provide information on dynamics and thermodynamics that generally is unavailable with cryo-EM alone. Accordingly, the synergetic use of HDX-MS and cryo-EM can validate each other and provide new insights[3]. We obtained a peptide coverage of 83% and 81% for prestin and SLC26A9, respectively, with a total of 266 and 338 peptides, allowing us to interrogate the protein-wide dynamics (**Fig. S5A; Table S1**). The main difference for the two proteins' sequence coverage is that only prestin has coverage at TM6 while only SLC26A9 has coverage at TM12. The different cleavage preferences at these two helices likely result from different flexiblity and/or exposure, which can be related to the different functions of the two proteins.

In 360 mM $Cl^-$, the majority of the TMDs for prestin and SLC26A9 had similar stability, with peptide-level protection factors (PFs) ranging from $10^3$ to $10^{6+}$; ΔG = 4.2 to 8.4+ kcal/mol (**Fig. S2 & S5B**). Both proteins had highly stable regions with negligible exchange after 27 h, our longest labeling time. We obtained near-residue level resolution at regions unresolved in the cryo-EM structures, including the intervening sequence of the STAS domain (sulfate transporter and anti-sigma factor antagonist) and the C-termini[2,4–7]. $\text{Regions}_{583\text{-}613,\ 734\text{-}764}$ for prestin and $\text{Regions}_{570\text{-}653,\ 741\text{-}770}$ for SLC26A9 had a PF of unity under all conditions examined (**Fig. S6**), indicating these regions are unfolded and independent of anion binding. This finding supports the proposal that the disordered regions of prestin may play a role in its interactions with other proteins for reasons of regulation rather than electromotility[8,9].

HDX is a solution-based method that probes the hydrogen bond network, and hence, any discrepancies with cryo-EM structures could reflect structural perturbations resulting from the sub-millisecond freezing process[10]. According to differences in cryo-EM and HDX, 2-3 residues form additional hydrogen bonds at the termini of two helices upon freezing. These sites included residues 565-566 and 720-722 for prestin and residues 225-226 and 738-740 for SLC26A9. Helical propagation can occur within 10 nsec[11] while the cooling time for cryo-EM can be as slow as 200 µsec[10], which provides ample time for helix extension as the helices equilibrate to the lower temperature[12]. Although we anticipate that such small-scale folding events are common during the freezing process, overall, the HDX data do not provide evidence for significant changes in hydrogen bonding patterns for either prestin or SLC26A9. We anticipate that both the barriers for larger-scale folding events and solvent viscosity increase significantly as the temperature drops, effectively trapping the protein in its pre-frozen conformation. Nevertheless, we note that motions that do not result in changes in the hydrogen bond network, such as rigid-body motions of helices, would not be identified by HDX.

### Supporting Information Text 3: Structure of prestin in HEPES and low $Cl^-$ levels.

Using single-particle cryo-EM, we set out to determine the structure of prestin in the HEPES-based buffer with the goal of visualizing a putative apo state. Prestin was initially screened in 190 mM HEPES in a nominal absence of $Cl^-$. This condition, however, led to widespread particle aggregation under cryogenic conditions. We reasoned that this aggregation may be linked to the already destabilized prestin without a bound $Cl^-$, as evidenced by HDX-MS data, as well as the low ionic strength of our buffer. Indeed, 1 mM $Cl^-$ sharply reduced particle aggregates, allowing us to solve the structure of prestin solubilized in GDN at a nominal resolution of 3.4 Å from particles, which corresponds to about 10% of the total particles in the sample. Surprisingly, under these conditions prestin adopted a “compact” conformation, virtually identical to the previously reported $Cl^-$-bound “Up” state (**Fig. S11**). Moreover, the anion-binding site is structurally indistinguishable from previous $Cl^-$-bound structures (**Fig. S11B**). Furthermore, when focusing on the anion-binding pocket in our cryo-EM map, we see clear evidence for an additional density, indicating that the pocket is occupied by a substrate (**Fig. S11C**). However, we were unable to model a HEPES anion into the binding pocket without substantial steric clashes. We therefore suggest that the resolved density represents instead a $Cl^-$ anion, given that a small population of $Cl^-$-bound prestin will be present from a weak $Cl^-$ affinity (e.g., $EC_{50}$=6 mM[13] implies 17% bound). Although we cannot confirm this notion with absolute certainty, it is clear that our cryo-EM structure does not represent a true apo state of prestin and is consistent with the notion that unbound, apo prestin is conformationally unstable.

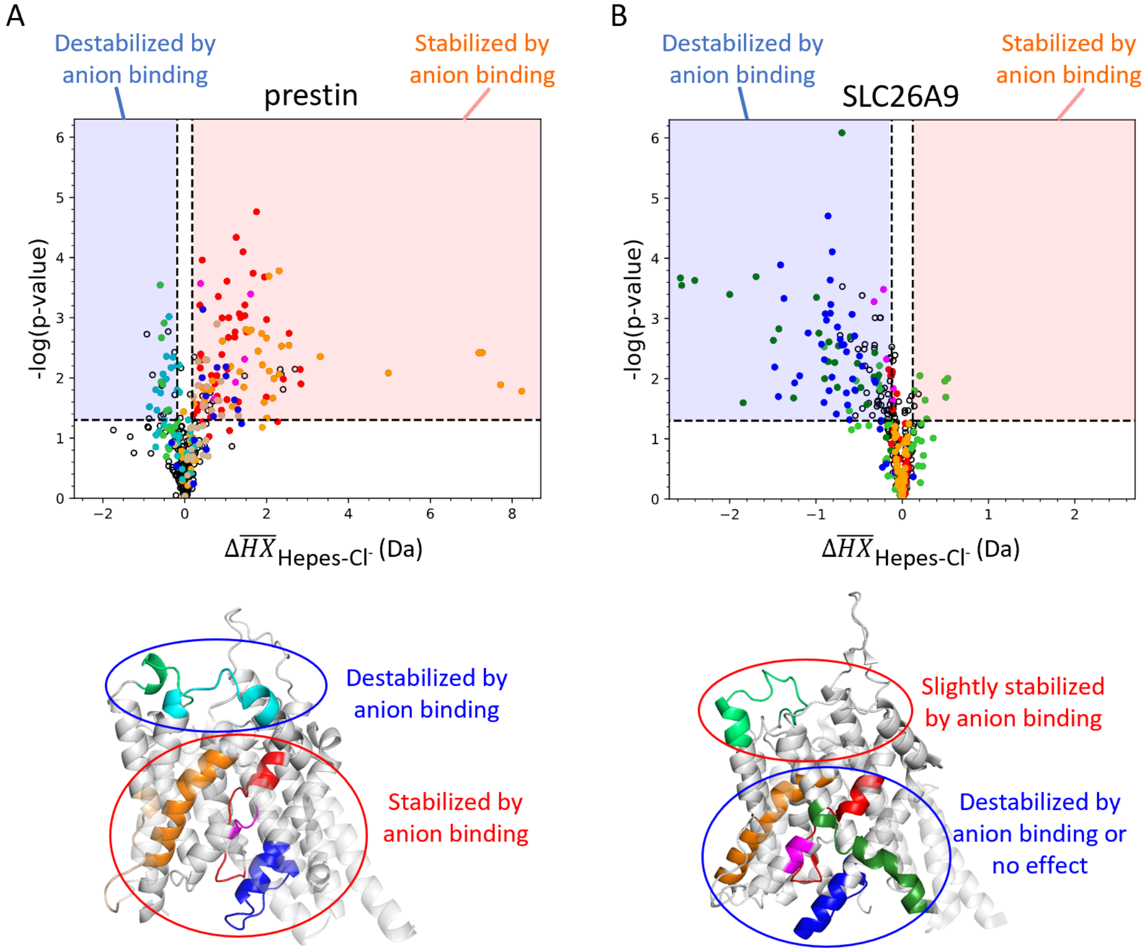


**Figure S1: Volcano plot analysis of HDX for prestin (A) and SLC26A9 (B) in response to $Cl^-$ binding.** A hybrid statistical analysis is employed[14]. Each data point represents one time point of a peptide. The horizontal axis shows the difference between the observed mean peptide masses ($\Delta\overline{HX}$) for the protein in HEPES and in $Cl^-$. The vertical axis shows Welch's t-test p-values. The dashed horizontal lines denote p-value significance limits defined at α = 0.05. The dashed vertical lines denote the $\Delta\overline{HX}$ significance threshold corresponding to α = 0.05, calculated to be ±0.17 Da and ±0.11 Da for **(A)** prestin and **(B)** SLC26A9, respectively. Data points in the red and blue area (i.e., exceeding significance limits in both dimensions) represent peptides that show statistical significance and are stabilized and destabilized upon $Cl^-$ binding, respectively. Colored data points correspond to colored regions in the corresponding TMD structure below.

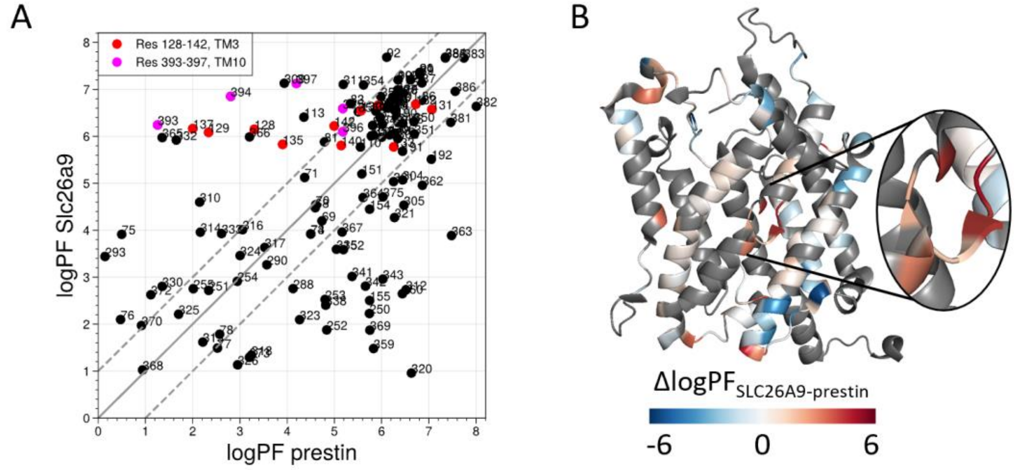


**Figure S2: Site-resolved protection factors for prestin and SLC26A9 obtained using PyHDX. (A)** Site-resolved protection factors in a log scale (logPF) for the TMDs of prestin and SLC26A9 with residue numbers denoted. **(B)** The difference in logPF (ΔlogPF) mapped onto the prestin structure. Grey represents regions with no fitting data available. TM5 and TM12-14 do not have fitting data and are not shown to highlight the anion-binding site. Fitting was done using PyHDX[15] on HDX-MS data measured for both proteins in $Cl^-$.

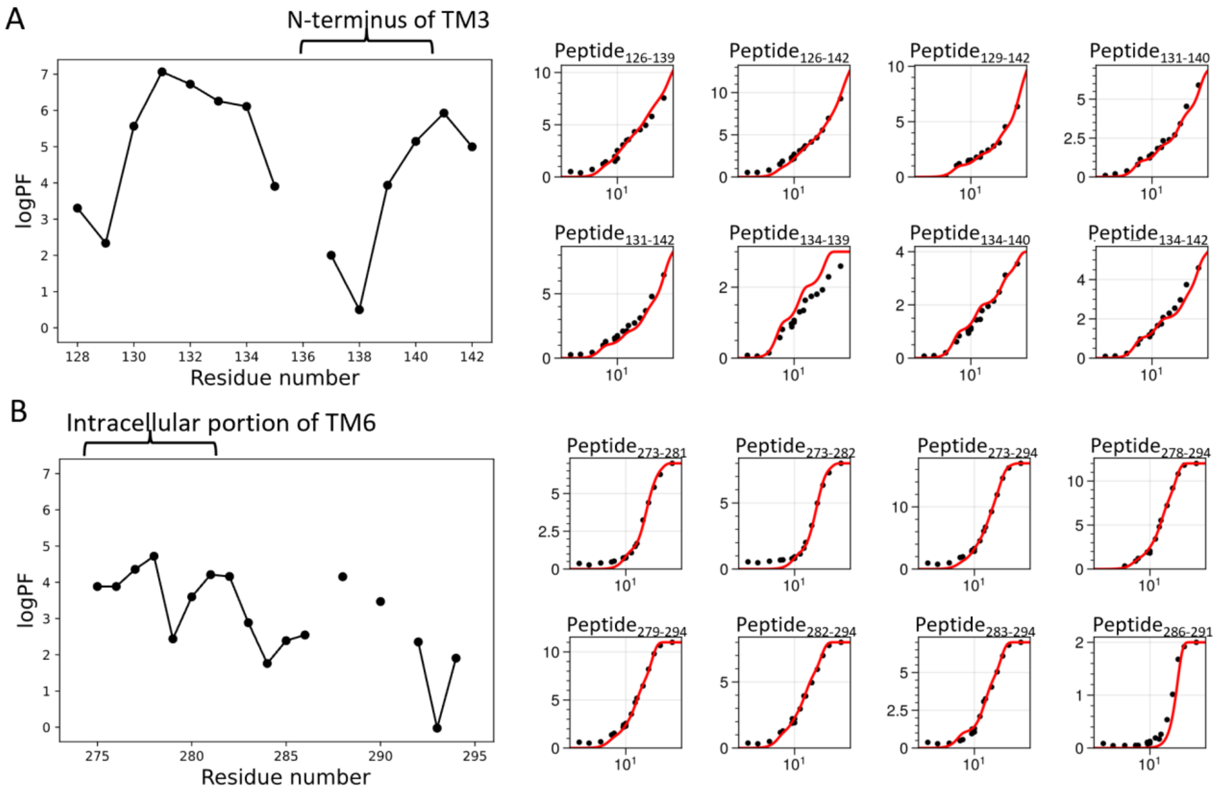


**Figure S3: PyHDX fitting supports that prestin exhibits helix fraying at the N-terminus of TM3 and mild cooperativity at the intracellular portion of TM6. Left:** Site-resolved protection factor values in a log scale (logPF) obtained by deconvoluting HDX-MS data using PyHDX[15] for regions covering **(A)** the N-terminus of TM3 and **(B)** the intracellular portion of TM6. **Right:** PyHDX fittings for the corresponding peptides – black circles: experimental deuteration levels; red curves: fittings.

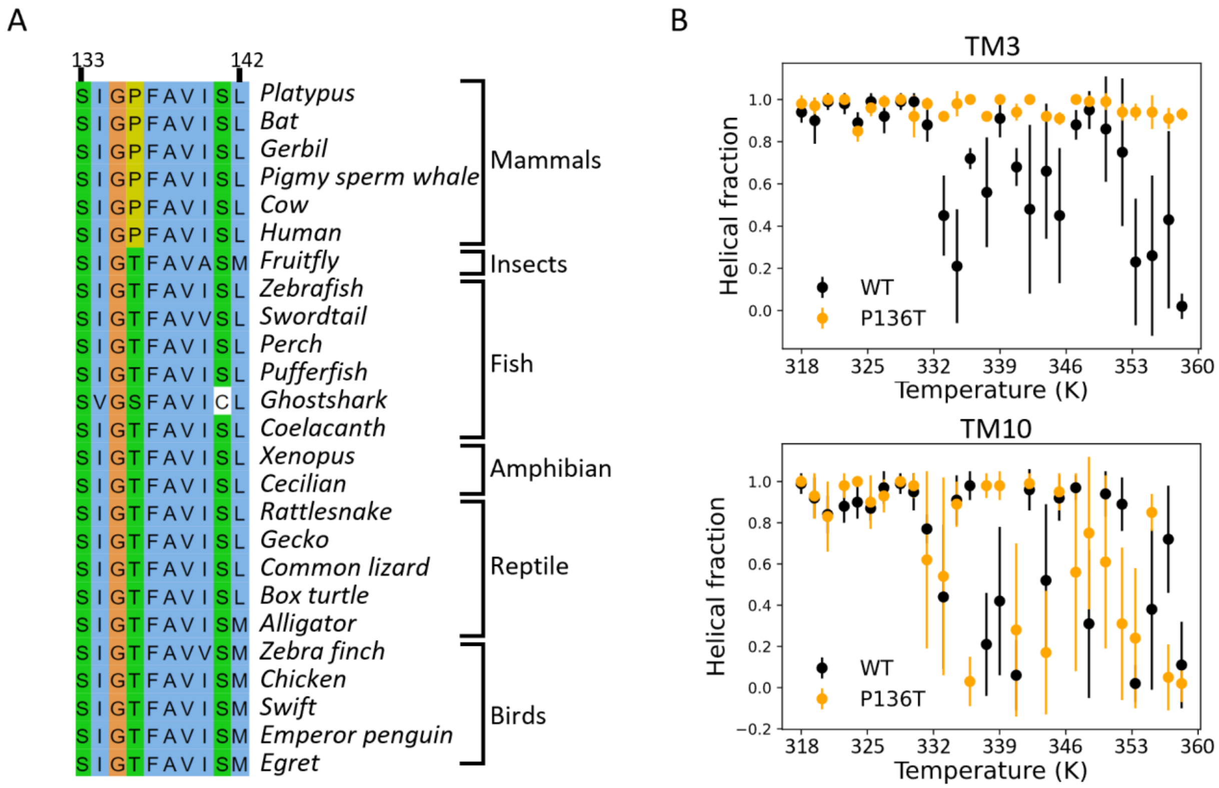


**Figure S4: Mammalian prestin has a conserved and helix-destabilizing proline 136 on TM3. (A)** Sequence alignment for the N-terminus of TM3 for prestin across species. A conserved threonine at residue 136 in prestin from other vertebrates is replaced with a proline in mammalian prestin. **(B)** Molecular simulations of thermo-denaturation of prestin conducted with Upside[16,17] (**Materials and Methods**). A P136T mutation largely increases the stability of TM3 but mildly decreases TM10 stability, as compared to wild-type prestin.

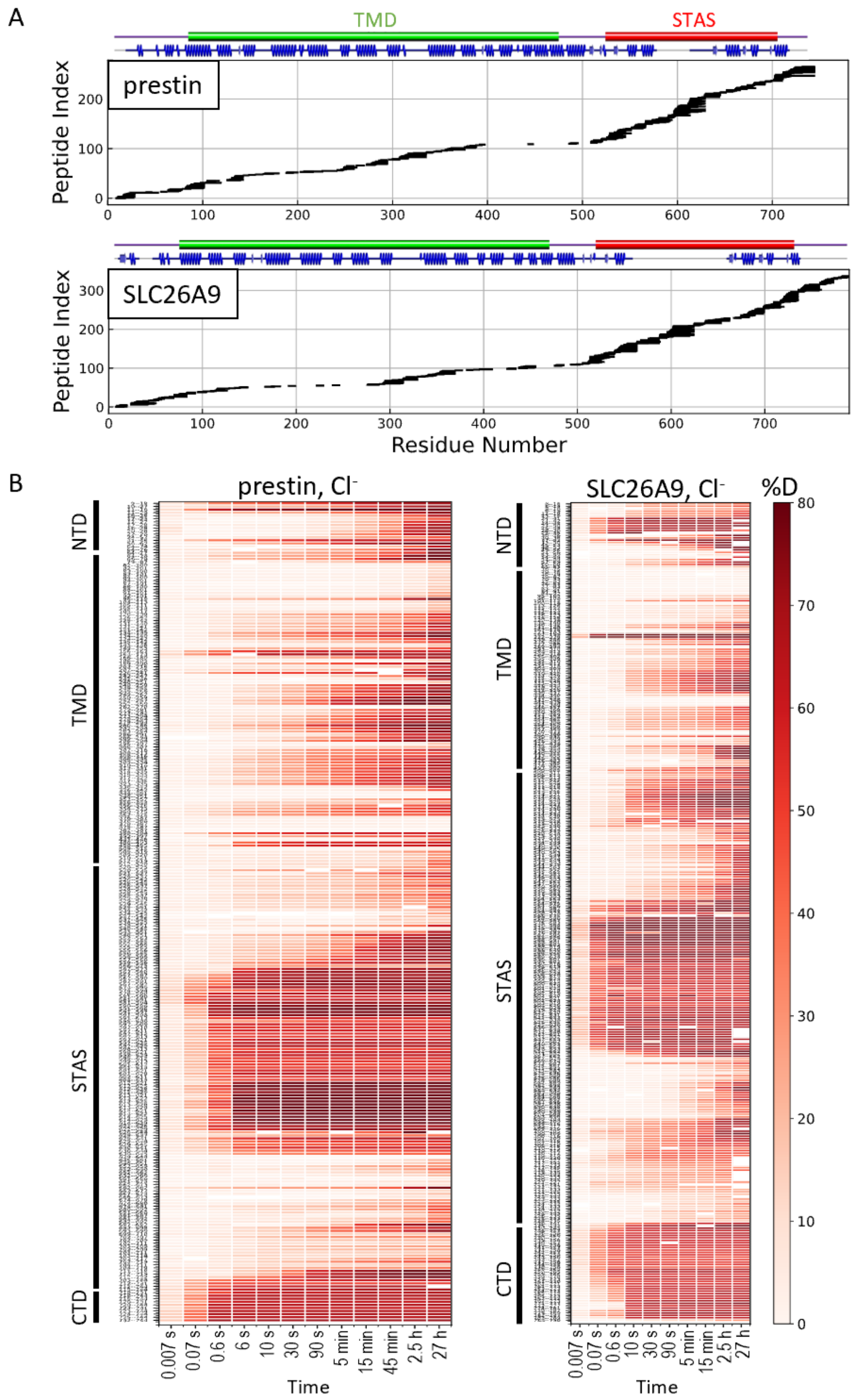


**Figure S5: HDX-MS sequence coverage and measurements for prestin and SLC26A9 in $Cl^-$. (A)** Peptide sequence coverage for prestin and SLC26A9 suitable for HDX-MS analysis. On the top indicates the sequence boundary for domains and secondary structures. **(B)** Heatmaps showing deuteration levels of all the peptides at each labeling time for prestin and SLC26A9 measured in $Cl^-$. Peptide sequences are displayed on the y-axis and legible through high resolution images.

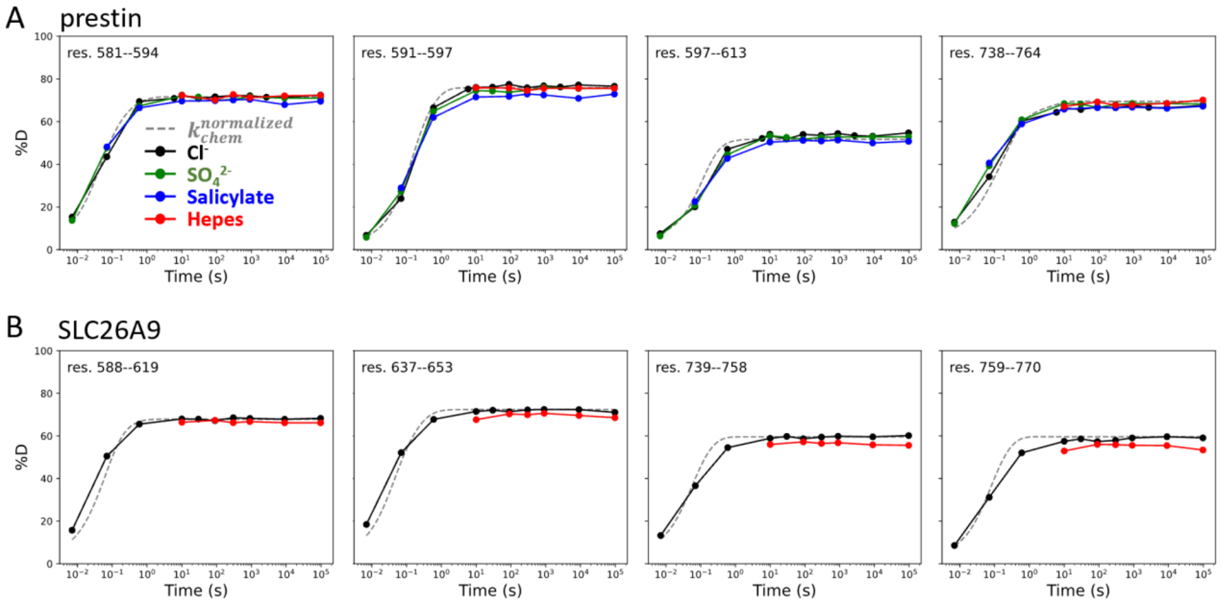


**Figure S6: Regions unresolved in cryo-EM structures are unfolded in all conditions examined.** Deuterium uptake plots for example peptides covering regions unresolved in cryo-EM structures for **(A)** prestin and **(B)** SLC26A9: black, $Cl^-$; green: $SO_4^{2-}$; blue: salicylate; red: HEPES. Grey dashed curves represent deuterium uptake with $k_{chem}$, normalized with the in- and back-exchange levels. Only one replicate for $Cl^-$ and HEPES are shown for clarity.

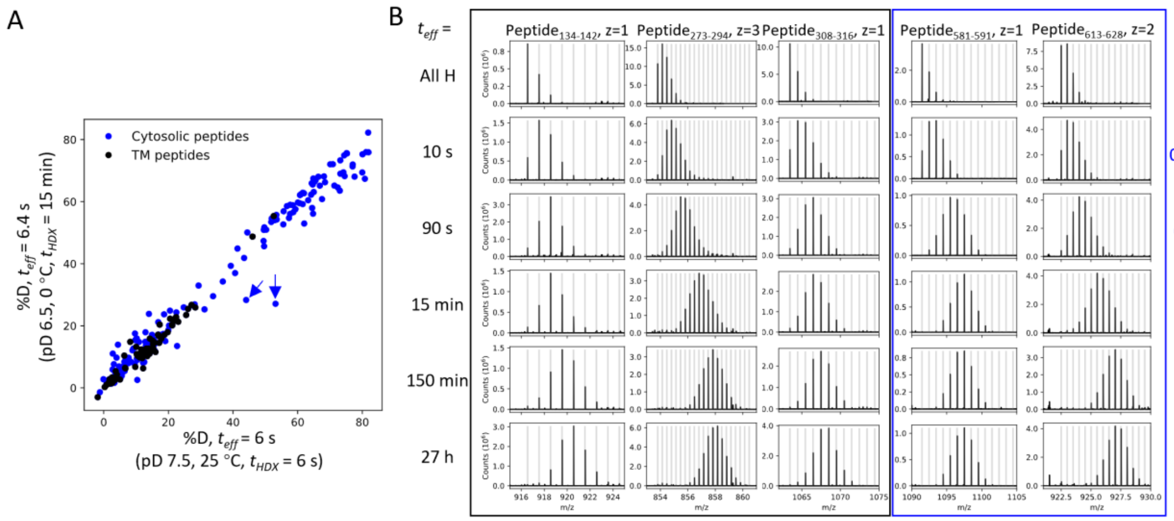


**Figure S7: HDX for prestin occurs via EX2 mechanism. (A)** Comparison of deuteration levels of all prestin peptides labeled under two HDX conditions: 1) $pD_{read}$ 7.1, 25 °C, $t_{HDX}$ = 6 s; 2) $pD_{read}$ 6.1, 0 °C, $t_{HDX}$ = 15 min, $t_{eff}$ = 6.4 s, where $t_{eff}$ represents effective HDX labeling time in $pD_{read}$ 7.1, 25 °C (**Materials and Methods**). Black: transmembrane (TM) peptides, blue: cytosolic peptides. The two outliers denoted by the blue arrows represent peptides covering the Cα2 helix in the STAS domain, whose large apparent %D difference between the two conditions results from pH-/temperature-dependent dynamics (data not shown). **(B)** Mass spectra of a representative set of peptides showing progressive unimodal isotope envelopes towards high m/z over time. Grey horizontal bars indicate theoretical m/z values for the corresponding isotopes.

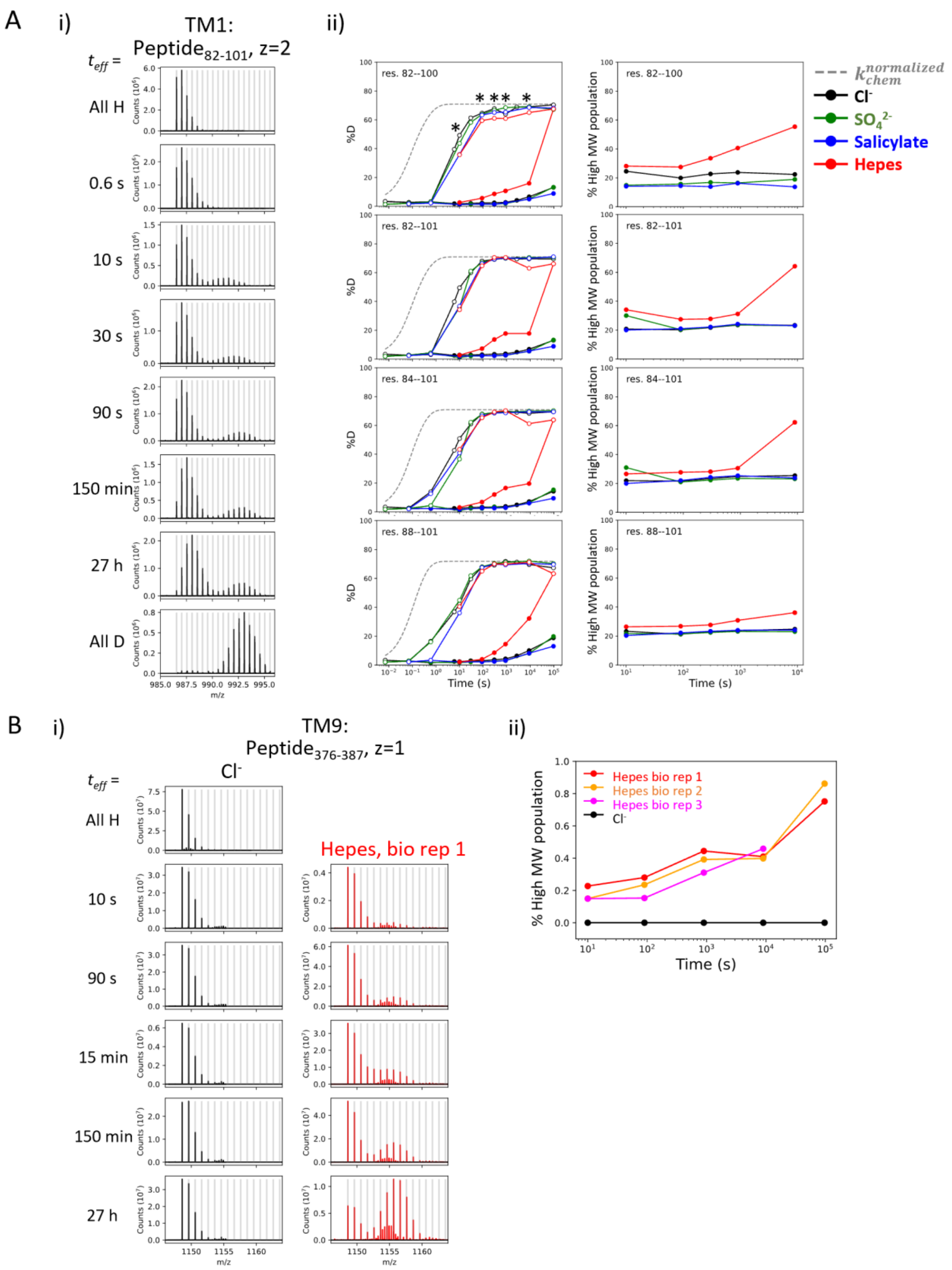


**Figure S8: Heterogeneity and HDX kinetics in TM1 and TM9. (A)** Conformational heterogeneity at TM1. **(i)** Example mass spectra for a TM1 peptide showing bimodal isotope envelopes, with both isotope distributions increasing in mass over time and exchanging via EX2 kinetics. **(ii) Left:** Deuterium uptake curves of example TM1 peptides plotting both the left (filled markers) and the right (empty markers) isotope envelopes under different anion conditions. Grey dashed curves represent deuterium uptake with $k_{chem}$, normalized with the in- and back-exchange levels. Only one replicate is shown for clarity. Asterisks represent time points used for the **(Right)** population fraction analysis, chosen as two isotope envelopes are well separated. Fractions of the heavier envelope (i.e., % High MW population) in HEPES are higher than those in other conditions as the left envelope merges into the right envelope, resulting in less distinct separation between the two envelopes. **(B)** HDX for prestin's TM9 exhibits EX1 kinetics in the apo state. **(i)** Example mass spectra for a TM9 peptide measured for prestin in $Cl^-$ (black) and HEPES (red). Identification of EX1 kinetics in HEPES is supported by the presence of two distinct mass envelopes, with the amplitude of the lighter envelope decreasing with a commensurate increase in the heavier envelope over time[1]. **(ii)** The fraction of the heavier envelope over time for prestin in $Cl^-$ (black) and HEPES (3 biological replicates are shown).

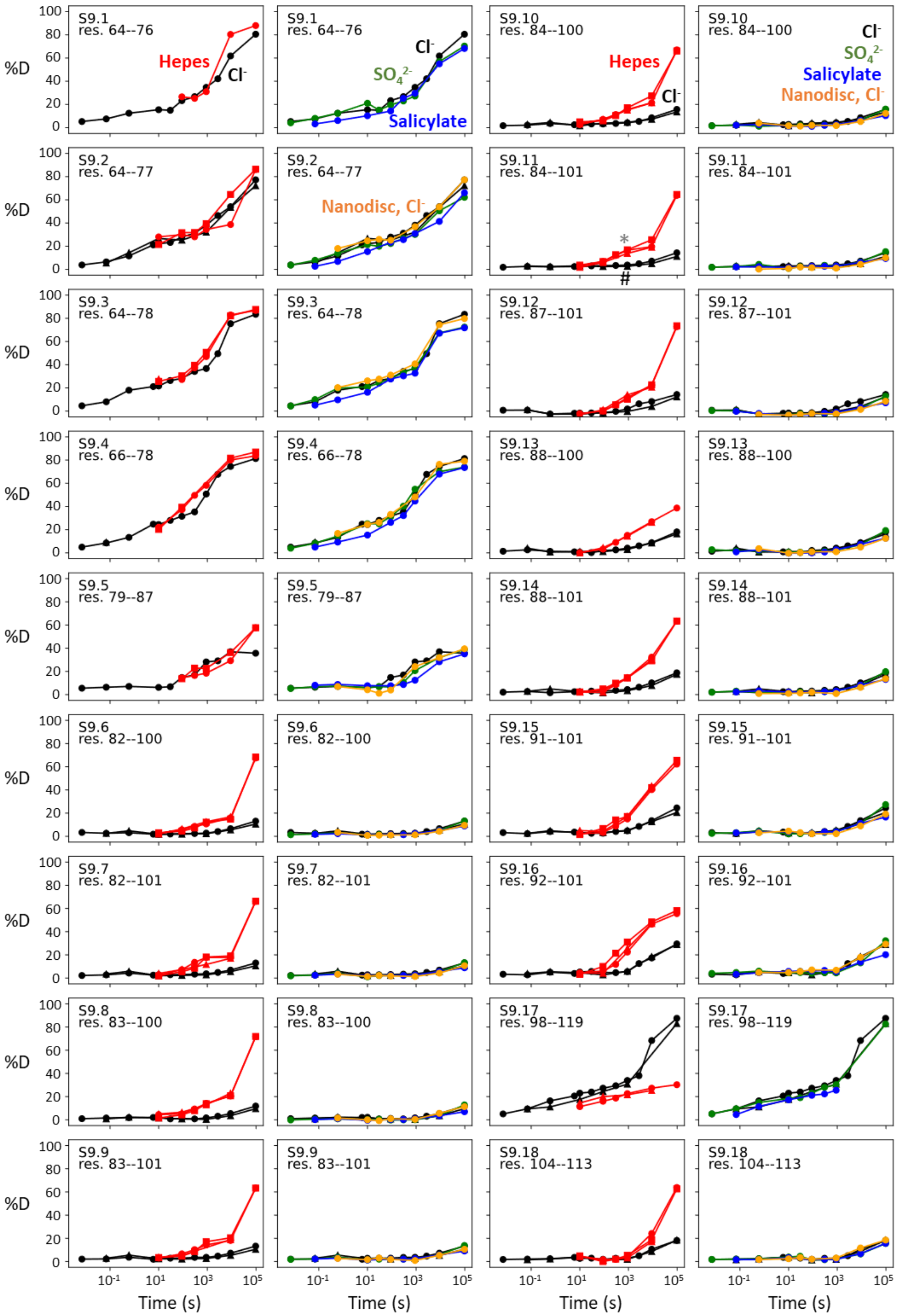
S9.1 res. 64--76
Hepes
Cl-
S9.1 res. 64--76
Cl-
SO4 2-
Salicylate
S9.10 res. 84--100
Hepes
Cl-
S9.10 res. 84--100
Cl-
SO4 2-
Salicylate
Nanodisc, Cl-
S9.2 res. 64--77
S9.2 res. 64--77
Nanodisc, Cl-
S9.11 res. 84--101
*
#
S9.11 res. 84--101
S9.3 res. 64--78
S9.3 res. 64--78
S9.12 res. 87--101
S9.12 res. 87--101
S9.4 res. 66--78
S9.4 res. 66--78
S9.13 res. 88--100
S9.13 res. 88--100
S9.5 res. 79--87
S9.5 res. 79--87
S9.14 res. 88--101
S9.14 res. 88--101
S9.6 res. 82--100
S9.6 res. 82--100
S9.15 res. 91--101
S9.15 res. 91--101
S9.7 res. 82--101
S9.7 res. 82--101
S9.16 res. 92--101
S9.16 res. 92--101
S9.8 res. 83--100
S9.8 res. 83--100
S9.17 res. 98--119
S9.17 res. 98--119
S9.9 res. 83--101
S9.9 res. 83--101
S9.18 res. 104--113
S9.18 res. 104--113
%D
Time (s)

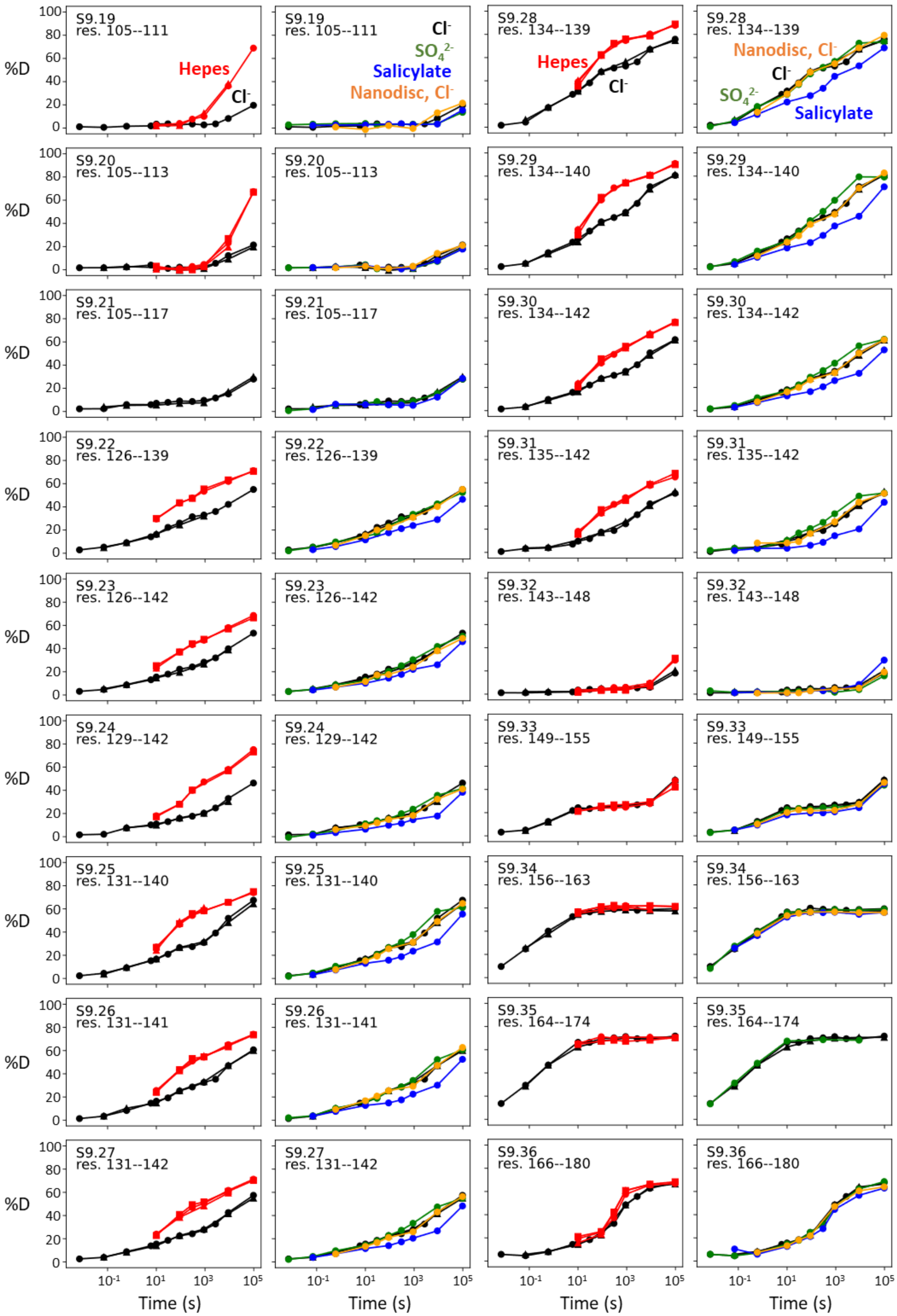
S9.19 res. 105--111
Hepes
Cl-
SO42-
Salicylate
Nanodisc, Cl-
S9.28 res. 134--139
S9.20 res. 105--113
S9.29 res. 134--140
S9.21 res. 105--117
S9.30 res. 134--142
S9.22 res. 126--139
S9.31 res. 135--142
S9.23 res. 126--142
S9.32 res. 143--148
S9.24 res. 129--142
S9.33 res. 149--155
S9.25 res. 131--140
S9.34 res. 156--163
S9.26 res. 131--141
S9.35 res. 164--174
S9.27 res. 131--142
S9.36 res. 166--180
%D
Time (s)

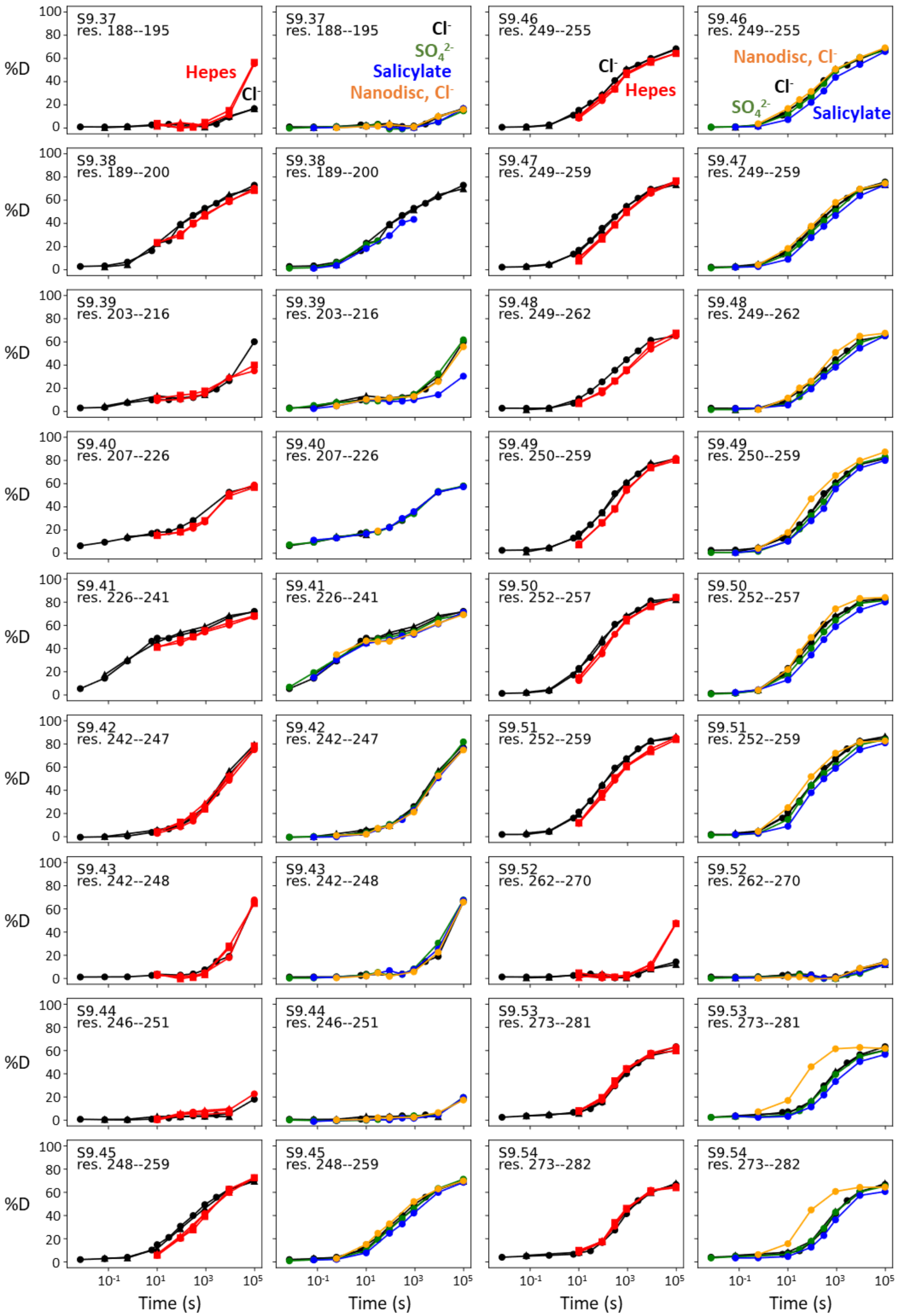
S9.37
res. 188--195
Hepes
Cl-
S9.37
res. 188--195
Cl-
SO4 2-
Salicylate
Nanodisc, Cl-
S9.46
res. 249--255
Cl-
Hepes
S9.46
res. 249--255
Nanodisc, Cl-
Cl-
SO4 2-
Salicylate
S9.38
res. 189--200
S9.38
res. 189--200
S9.47
res. 249--259
S9.47
res. 249--259
S9.39
res. 203--216
S9.39
res. 203--216
S9.48
res. 249--262
S9.48
res. 249--262
S9.40
res. 207--226
S9.40
res. 207--226
S9.49
res. 250--259
S9.49
res. 250--259
S9.41
res. 226--241
S9.41
res. 226--241
S9.50
res. 252--257
S9.50
res. 252--257
S9.42
res. 242--247
S9.42
res. 242--247
S9.51
res. 252--259
S9.51
res. 252--259
S9.43
res. 242--248
S9.43
res. 242--248
S9.52
res. 262--270
S9.52
res. 262--270
S9.44
res. 246--251
S9.44
res. 246--251
S9.53
res. 273--281
S9.53
res. 273--281
S9.45
res. 248--259
S9.45
res. 248--259
S9.54
res. 273--282
S9.54
res. 273--282
%D
0
20
40
60
80
100
10-1
101
103
105
Time (s)

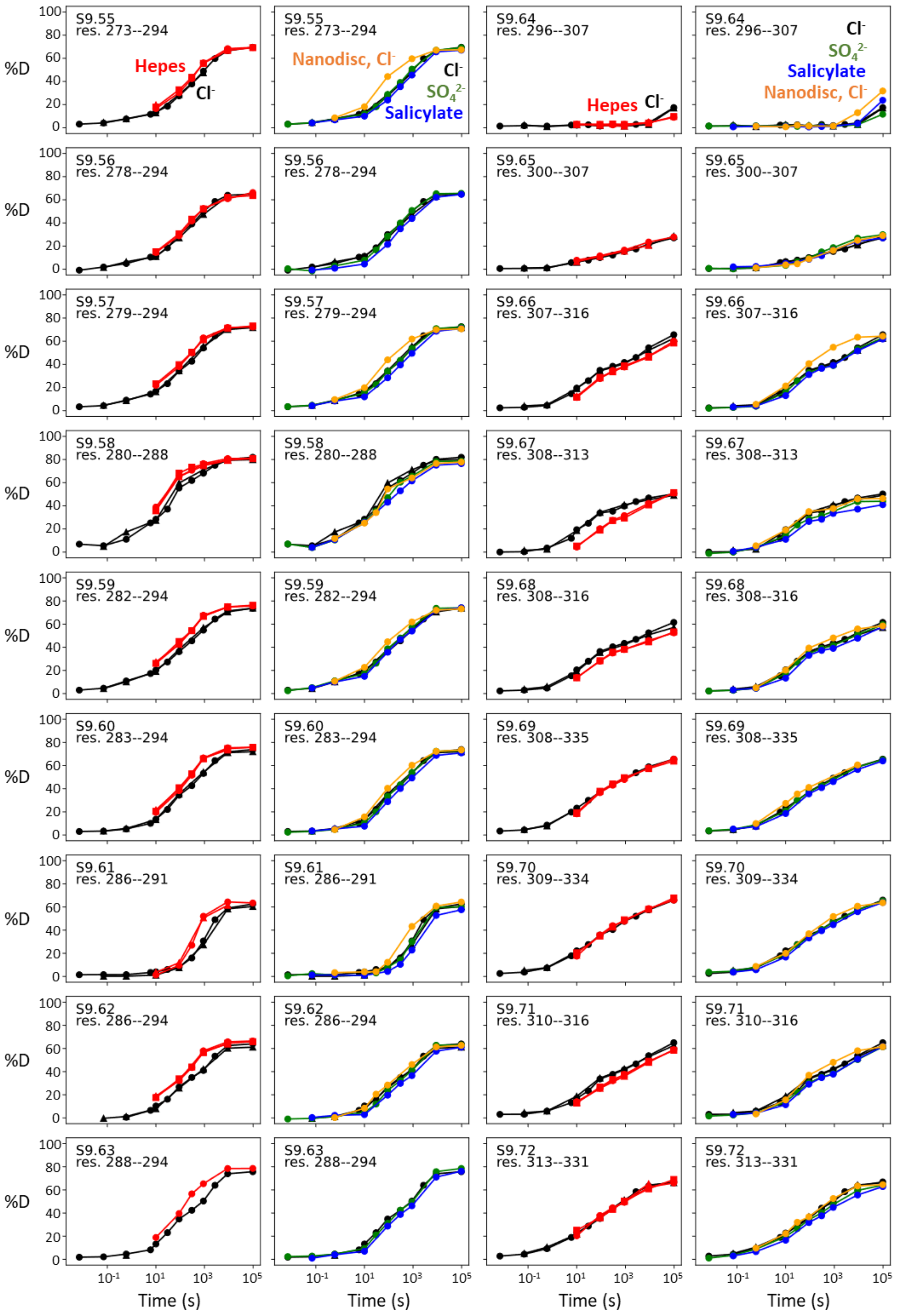

S9.55 res. 273--294
Hepes
Cl⁻
Nanodisc, Cl⁻
SO4 2-
Salicylate
S9.64 res. 296--307
S9.56 res. 278--294
S9.65 res. 300--307
S9.57 res. 279--294
S9.66 res. 307--316
S9.58 res. 280--288
S9.67 res. 308--313
S9.59 res. 282--294
S9.68 res. 308--316
S9.60 res. 283--294
S9.69 res. 308--335
S9.61 res. 286--291
S9.70 res. 309--334
S9.62 res. 286--294
S9.71 res. 310--316
S9.63 res. 288--294
S9.72 res. 313--331
%D
Time (s)

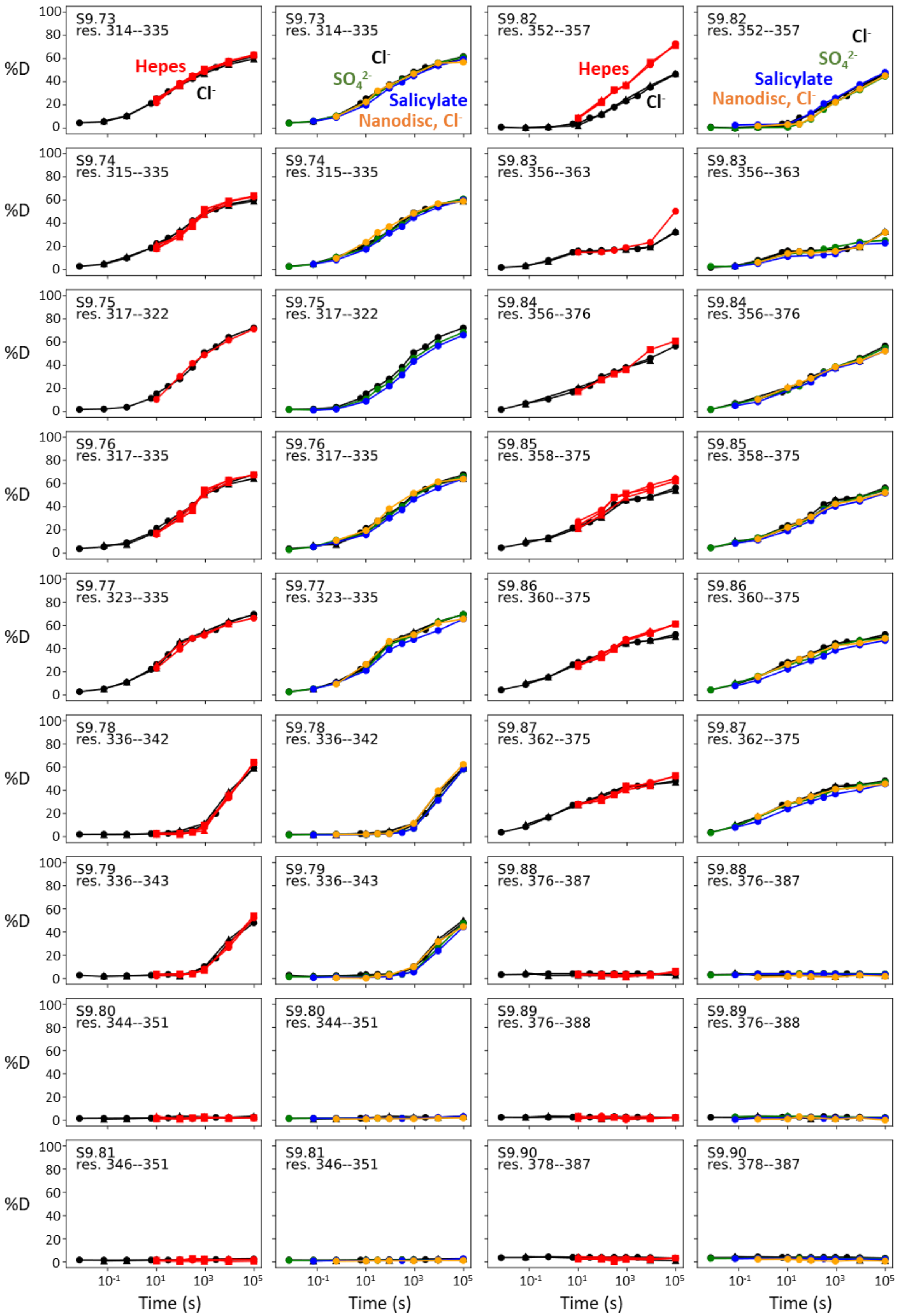
S9.73 res. 314--335
Hepes
Cl-
SO4 2-
Salicylate
Nanodisc, Cl-
S9.82 res. 352--357
S9.74 res. 315--335
S9.83 res. 356--363
S9.75 res. 317--322
S9.84 res. 356--376
S9.76 res. 317--335
S9.85 res. 358--375
S9.77 res. 323--335
S9.86 res. 360--375
S9.78 res. 336--342
S9.87 res. 362--375
S9.79 res. 336--343
S9.88 res. 376--387
S9.80 res. 344--351
S9.89 res. 376--388
S9.81 res. 346--351
S9.90 res. 378--387
%D
Time (s)

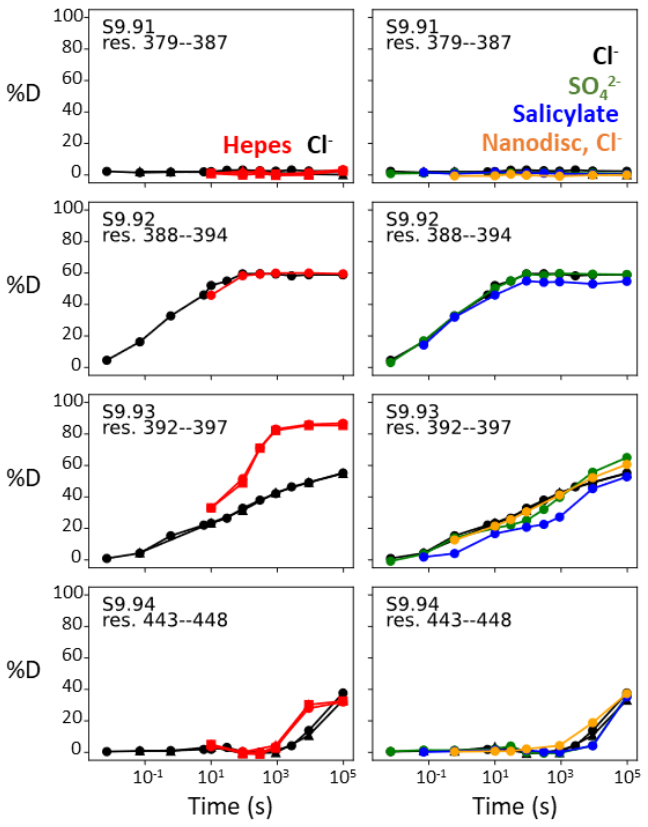


**Figure S9: Deuterium uptake curves for all peptides covering prestin's transmembrane domain.** Two deuterium uptake plots are shown for each peptide – left plot: $Cl^-$ (black) and HEPES (red); right plot: $Cl^-$ (black), $SO_4^{2-}$ (green), salicylate (blue), and prestin in nanodisc in $Cl^-$ (orange). Except for the orange curves, all other HDX data were collected on GDN-solublized prestin. Replicates (circles, triangles, and squares): 2 in $Cl^-$, 3 in HEPES, biological. Peptides covering TM9 (S9.88-S9.91) exhibited EX1 behavior in HEPES (**Fig. S8B**).

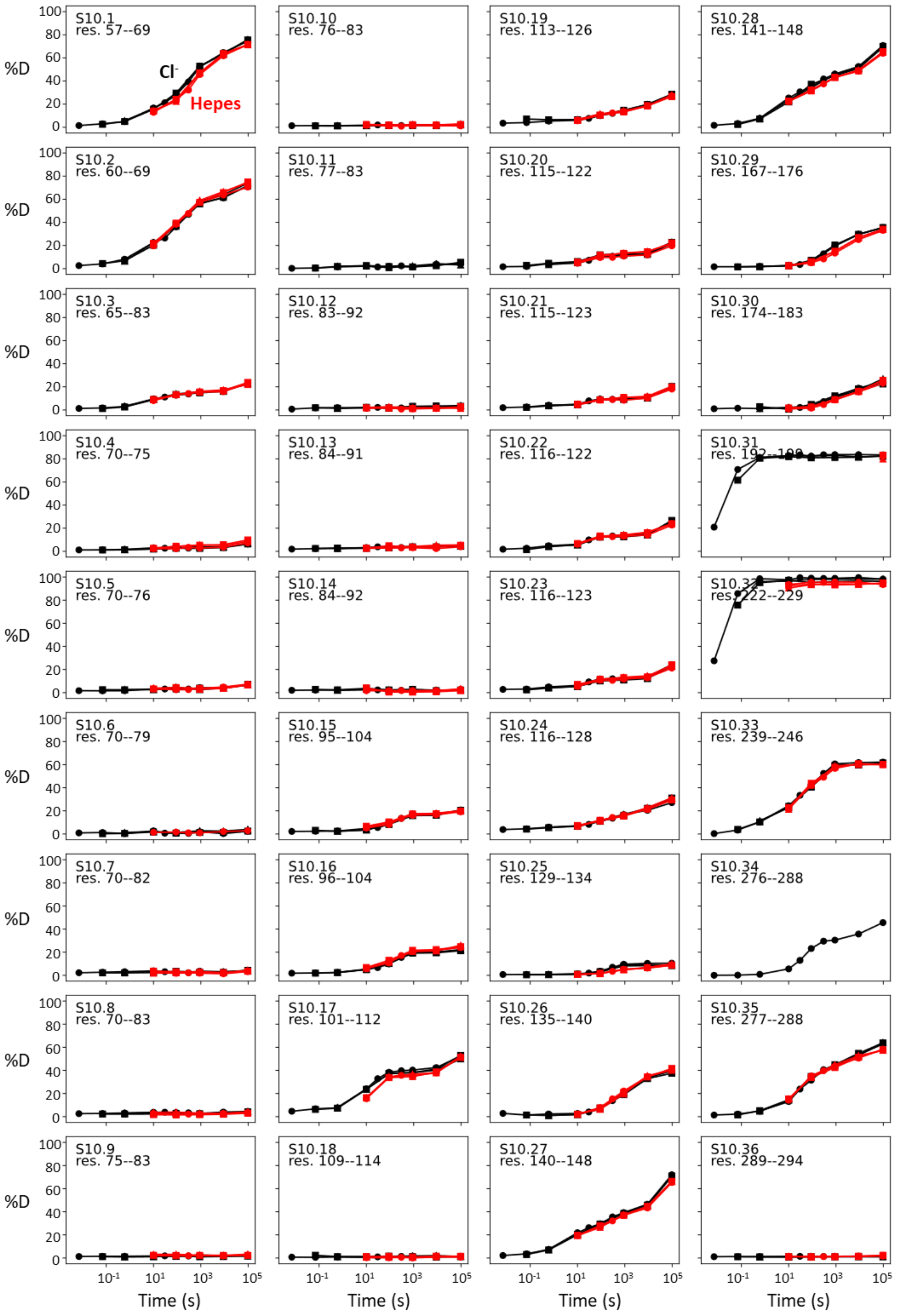
%D
Time (s)
Cl-
Hepes
S10.1 res. 57--69
S10.2 res. 60--69
S10.3 res. 65--83
S10.4 res. 70--75
S10.5 res. 70--76
S10.6 res. 70--79
S10.7 res. 70--82
S10.8 res. 70--83
S10.9 res. 75--83
S10.10 res. 76--83
S10.11 res. 77--83
S10.12 res. 83--92
S10.13 res. 84--91
S10.14 res. 84--92
S10.15 res. 95--104
S10.16 res. 96--104
S10.17 res. 101--112
S10.18 res. 109--114
S10.19 res. 113--126
S10.20 res. 115--122
S10.21 res. 115--123
S10.22 res. 116--122
S10.23 res. 116--123
S10.24 res. 116--128
S10.25 res. 129--134
S10.26 res. 135--140
S10.27 res. 140--148
S10.28 res. 141--148
S10.29 res. 167--176
S10.30 res. 174--183
S10.31
S10.32 res. 222--229
S10.33 res. 239--246
S10.34 res. 276--288
S10.35 res. 277--288
S10.36 res. 289--294

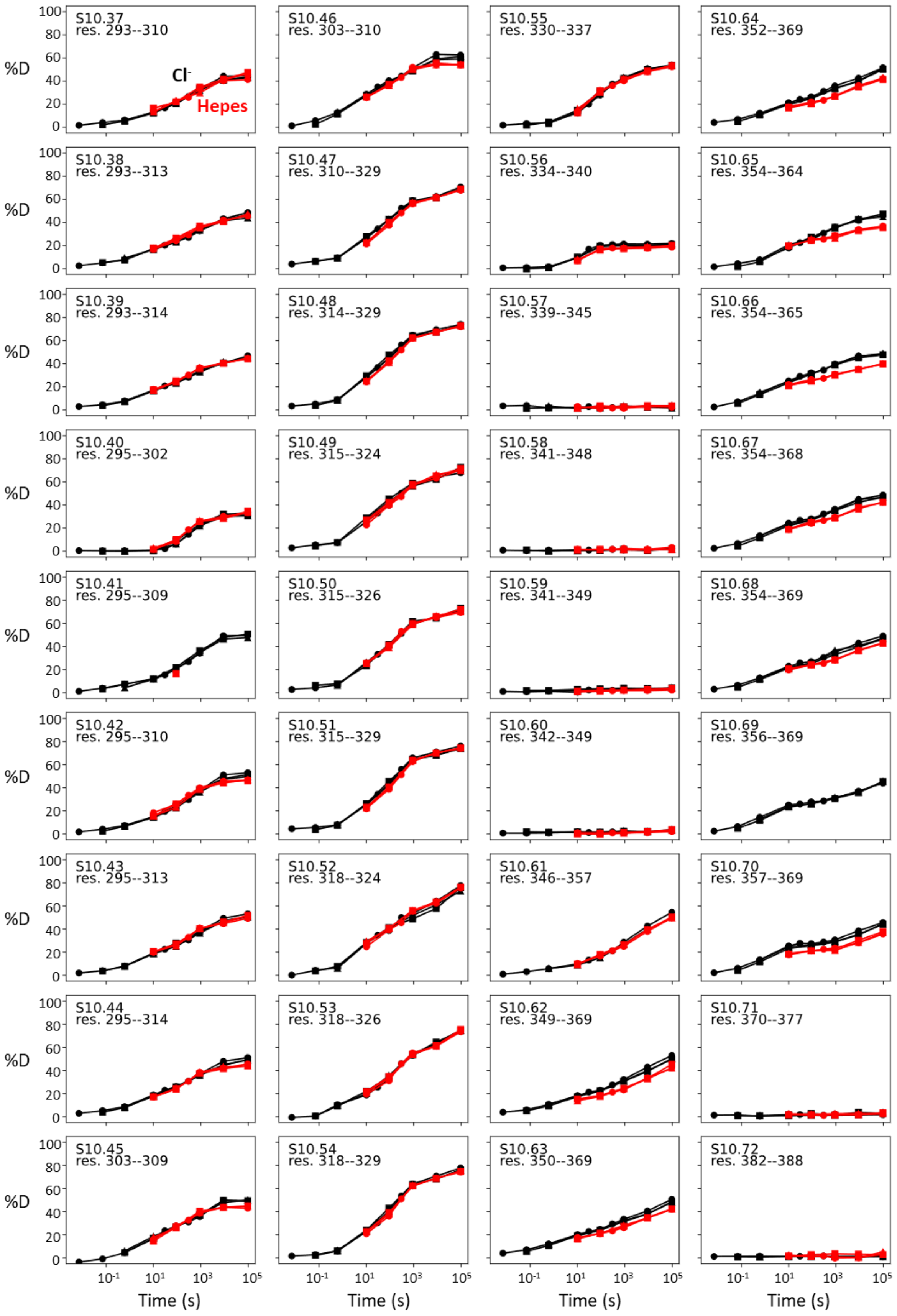
S10.37 res. 293--310
Cl-
Hepes
S10.38 res. 293--313
S10.39 res. 293--314
S10.40 res. 295--302
S10.41 res. 295--309
S10.42 res. 295--310
S10.43 res. 295--313
S10.44 res. 295--314
S10.45 res. 303--309
S10.46 res. 303--310
S10.47 res. 310--329
S10.48 res. 314--329
S10.49 res. 315--324
S10.50 res. 315--326
S10.51 res. 315--329
S10.52 res. 318--324
S10.53 res. 318--326
S10.54 res. 318--329
S10.55 res. 330--337
S10.56 res. 334--340
S10.57 res. 339--345
S10.58 res. 341--348
S10.59 res. 341--349
S10.60 res. 342--349
S10.61 res. 346--357
S10.62 res. 349--369
S10.63 res. 350--369
S10.64 res. 352--369
S10.65 res. 354--364
S10.66 res. 354--365
S10.67 res. 354--368
S10.68 res. 354--369
S10.69 res. 356--369
S10.70 res. 357--369
S10.71 res. 370--377
S10.72 res. 382--388
%D
Time (s)

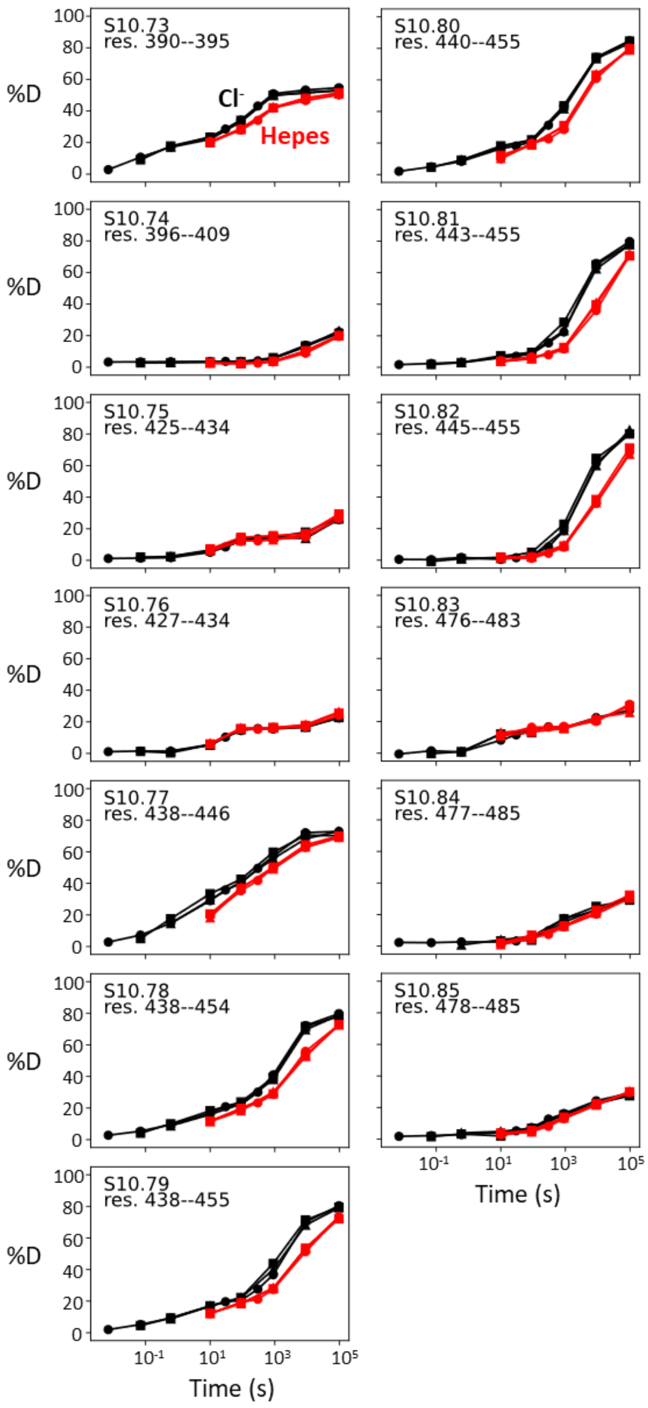


**Figure S10: Deuterium uptake curves for all peptides covering SLC26A9's transmembrane domain.** One deuterium uptake plot is shown for each peptide – $Cl^-$ (black) and HEPES (red). Three technical replicates are shown as circles, triangles, and squares. All HDX data were collected on GDN-solublized SLC26A9.

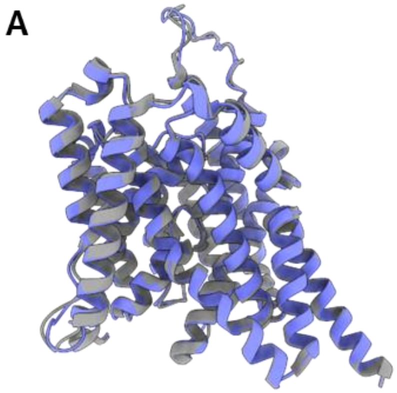


**Figure S11: The cryo-EM structure for prestin in a HEPES-based buffer containing 1 mM $Cl^-$ (blue; PDB 8UC1) highly resembles the structure in the reported $Cl^-$-bound state. (A)** Overlay of prestin's TMD with that solved in a high-chloride buffer (grey; PDB 7S8X). **(B)** Overlay of TM1, TM3, and TM10, with key residues that make up the anion-binding site. **(C)** Cryo-EM density forming the anion-binding site (blue). Additional density (red) that is incompatible with the placing of a HEPES molecule was resolved at the anion-binding site.

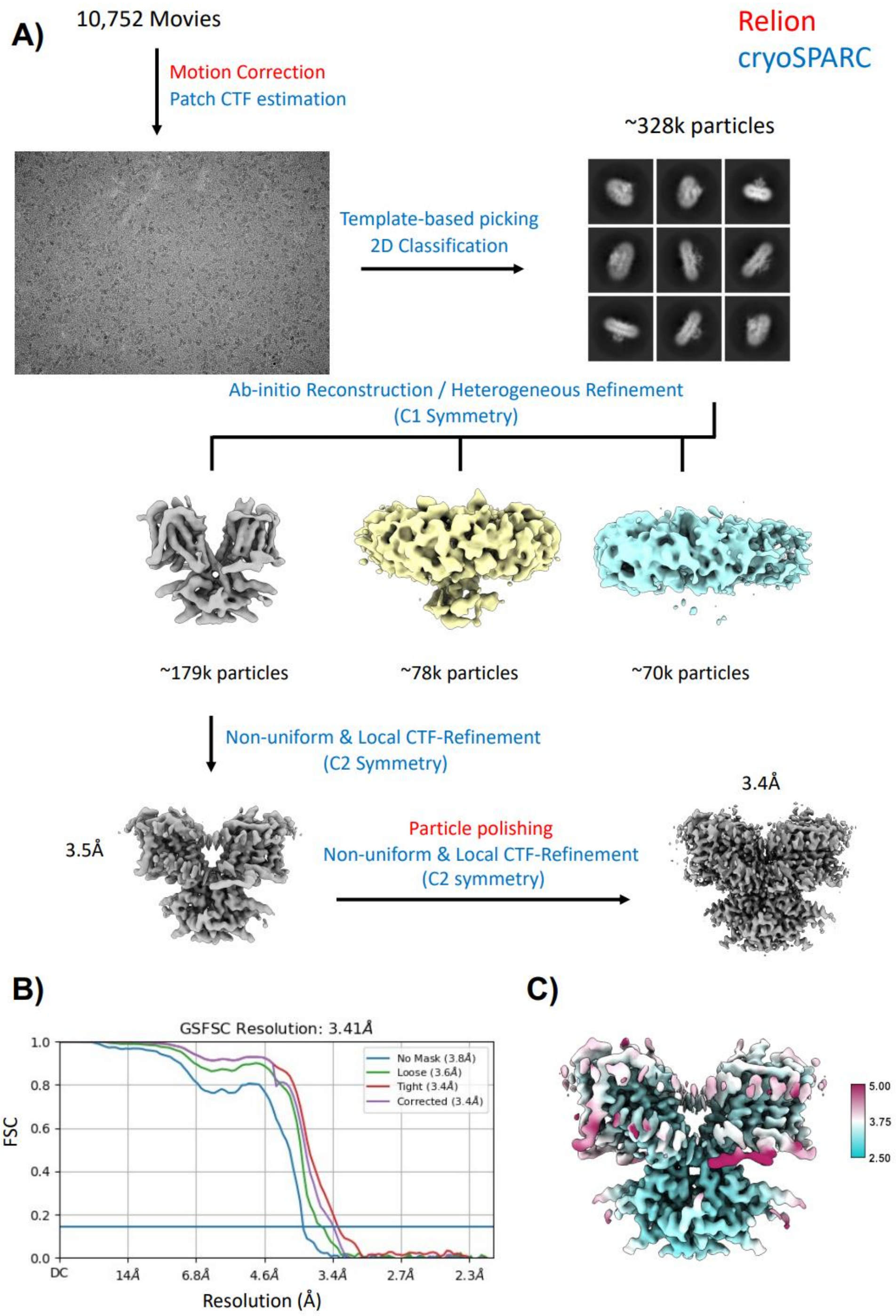


**Figure S12: Workflow for the processing of the cryo-EM data. (A)** Steps indicated in red font were performed in Relion, steps indicated in blue were performed in cryoSPARC. **(B)** FSC curve showing that the final reconstruction reached a nominal resolution of 3.4 Å (at FSC=0.143). **(C)** Local resolution estimation of the final reconstruction.

### Table S1: Biochemical and statistical details for HDX

| Dataset | Prestin, $Cl^-$ | Prestin, HEPES (apo) | prestin, $SO_4^{2-}$ | prestin, salicylate | prestin, nanodisc, $Cl^-$ | Slc26a9, $Cl^-$ | Slc26a9, HEPES (apo) |
|---|---|---|---|---|---|---|---|
| HDX reaction details | 360 mM NaCl, 20 mM Tris (NaPi for $pD_{read}$ 6.1, 0 °C), 3 mM DTT, 1 mM EDTA, 0.02% GDN. $pD_{read}$ 7.1, 25 °C or pD 6.5, 0 °C | 150 mM HEPES, 0.02% GDN. $pD_{read}$ 7.1, 25 °C | 140 mM $Na_2SO_4$, 5 mM $MgSO_4$, 20 mM NaPi, 0.02% GDN. $pD_{read}$ 7.1, 25 °C or $pD_{read}$ 6.1, 0 °C | 140 mM $Na_2SO_4$, 5 mM $MgSO_4$, 50 mM salicylate, 20 mM NaPi, 0.02% GDN. $pD_{read}$ 7.1, 25 °C or $pD_{read}$ 6.1, 0 °C | 20 mM Tris (NaPi for $pD_{read}$ 6.1, 0 °C), 150 mM NaCl. $pD_{read}$ 7.1, 25 °C or pD 6.5, 0 °C | 360 mM NaCl, 20 mM Tris (NaPi for pD 6.5, 0 °C), 3 mM DTT, 1 mM EDTA, 0.02% GDN. $pD_{read}$ 7.1, 25 °C or $pD_{read}$ 6.1, 0 °C | 150 mM HEPES, 0.02% GDN. $pD_{read}$ 7.1, 25 °C |
| HDX time course (*: replicated. Times in parenthesis: times in $pD_{read}$ 7.1, 25 °C after correcting for the $k_{chem}$ difference) | $pD_{read}$ 6.1, 0 °C: 1s (0.007s), 10s (0.07s)*, 90s (0.6s)*; $pD_{read}$ 7.1, 25 °C: 6s, 10s*, 30s, 90s*, 5min, 15min*, 45min, 150min*, 27h* | $pD_{read}$ 7.1, 25 °C: 10s*, 90s*, 5min*, 15min*, 150min*, 27h* | $pD_{read}$ 6.1, 0 °C: 1s (0.007s), 10s (0.07s), 90s (0.6s); $pD_{read}$ 7.1, 25 °C: 10s, 30s, 90s, 5min, 15min, 150min, 27h | $pD_{read}$ 6.1, 0 °C: 1s (0.007s), 90s (0.6s); $pD_{read}$ 7.1, 25 °C: 10s, 90s, 5min, 15min, 150min, 27h | $pD_{read}$ 6.1, 0 °C: 90s (0.6s); $pD_{read}$ 7.1, 25 °C: 10s, 90s, 15min, 150min, 27h | $pD_{read}$ 6.1, 0 °C: 1s (0.007s), 10s (0.07s)*, 90s (0.6s)*; $pD_{read}$ 7.1, 25 °C: 10s*, 30s, 90s*, 5min, 15min*, 150min*, 27h* | $pD_{read}$ 7.1, 25 °C: 10s*, 90s*, 15min*, 150min*, 27h* |
| HDX control samples | Non-deuterated control; in-exchange control; maximally labeled control | | | | | Non-deuterated control; in-exchange control; maximally labeled control | |

<table>
<tr><td>In- and back-exchange, mean/IQR</td><td colspan="5">In-exchange: 3.1% / 2.0%; back-exchange: 27% / 14%</td><td colspan="2">In-exchange: 2.5% / 1.9%; back-exchange: 29% / 17%</td></tr>
<tr><td>No. of peptides</td><td>266 (TMD: 95; cytosolic: 171)</td><td>265 (TMD: 94; cytosolic: 171)</td><td>266 (TMD: 95; cytosolic: 171)</td><td>265 (TMD: 94; cytosolic: 171)</td><td>256 (TMD: 85; cytosolic: 171)</td><td>338 (TMD: 85; cytosolic: 253)</td><td>335 (TMD: 82; cytosolic: 253)</td></tr>
<tr><td>Sequence coverage</td><td>83% (TMD: 75%; cytosolic: 95%)</td><td>83% (TMD: 75%; cytosolic: 95%)</td><td>83% (TMD: 75%; cytosolic: 95%)</td><td>83% (TMD: 74%; cytosolic: 95%)</td><td>79% (TMD: 67%; cytosolic: 95%)</td><td>81% (TMD: 68%; cytosolic: 96%)</td><td>81% (TMD: 68%; cytosolic: 96%)</td></tr>
<tr><td>Average peptide length/redundancy</td><td>12.2/4.3</td><td>12.2/4.3</td><td>12.2/4.3</td><td>12.2/4.3</td><td>12.2/4.1</td><td>12.5/5.3</td><td>12.5/5.3</td></tr>
<tr><td>Replicates</td><td>2 (Biological)</td><td>3 (Biological)</td><td>1</td><td>1</td><td>1</td><td>3 (Technical)</td><td>3 (Technical)</td></tr>
<tr><td>Repeatability (TM peptides only)</td><td>0.69%/0.06 Da (average SD of the Δ%D/Δ#D between the duplicates)</td><td>0.93%/0.09 Da (average SD)</td><td>N/A</td><td>N/A</td><td>N/A</td><td>0.64%/0.06 Da (average SD)</td><td>0.60%/0.05 Da (average SD)</td></tr>
<tr><td rowspan="2">Significant differences in HDX (ΔHDX > X Da, TM peptides only)</td><td>N/A</td><td>0.22 Da (95% CI)</td><td>N/A</td><td>N/A</td><td>N/A</td><td>0.14 Da (95% CI)</td><td>0.13 Da (95% CI)</td></tr>
<tr><td colspan="2">0.17 Da (95% CI)</td><td>N/A</td><td>N/A</td><td>N/A</td><td colspan="2">0.11 Da (95% CI)</td></tr>
</table>
